# Ancient pangenomic origins of noncanonical NLR genes underlying the recent evolutionary rescue of a staple crop

**DOI:** 10.1101/2025.04.11.648396

**Authors:** Carl J. VanGessel, Terry J. Felderhoff, Daniil M. Prigozhin, Meihua Cui, Gael Pressoir, Adam L. Healey, John T. Lovell, Vamsi J. Nalam, Marc T. Nishimura, Geoffrey P. Morris

## Abstract

Evolutionary rescue occurs when populations in deteriorating environments avoid extinction by rapid adaptation. The recent evolutionary rescue of the cereal crop sorghum via *RMES1* aphid resistance is among a few known in situ cases, but its pangenomic origins and molecular basis is not yet known. Here, we describe the behavioral effects, molecular endophenotypes, and pangenomic evolution underlying this evolutionary rescue. Analysis of near-isogenic lines show that *RMES1* disrupts phloem feeding via global immunity activation of conserved defense networks. A growth-to-defense transition is evidenced by extensive transcriptome remodeling (>15% of expressed genes) and mediated by salicylic acid signaling. Nucleotide-binding leucine-rich repeat (NLR) immune receptor genes at the *RMES1* locus harbored on a large tandem duplication have extensive copy number variation across the sorghum pangenome. The likely causative NLRs (RMES1A and RMES1B) were inferred from expression and structural variation. The NLRs encoded at RMES1 lack an N-terminal signaling domain and have nucleotide-binding domain sequence variation expected to result in a loss of ATP binding, suggesting RMES1 NLRs function via a noncanonical mechanism. The *RMES1* NLR family is shared across the grass super-pangenome and includes the brown planthopper resistance gene *BPH40* in *Oryza sativa*, which is syntenic to *RMES1*. Finally, k-mer analysis of *RMES1* haplotypes in the sorghum pangenome established the East African origin of rare standing variation for resistance. Thus, the birth-and-death process at an ancient gene cluster generated pangenomic variation that was recruited to activate coordinated defense pathways and provide evolutionary rescue.

## INTRODUCTION

Global change is causing environmental degradation impacting ecosystems (*1*). Ecological adaptation in response to these disturbances occurs via diverse genetic and molecular mechanisms (*2*). Evolutionary rescue is a form of adaptation that prevents extinction or extirpation in response to an acute stress (*3*, *4*). Evolutionary rescue is expected to be driven by Mendelian traits encoded by single genes that undergo a selective sweep whereas oligogenic or polygenic architecture may not be able to fix resilience quickly enough (*5*). In plants, molecular machinery responsible for evolutionary rescue has been described for flowering time pathways (*6*, *7*). Conserved pathways such as photoperiodism can be modulated for evolutionary rescue by loss of function of canonical flowering time genes (*7*). However, few studies have examined how evolutionary rescue manifests in the context of plant resistance to insect herbivores, particularly through immune receptor-mediated defense responses. Investigating the biological components of evolutionary rescue in tandem with simulations is necessary to address global environmental change. The recent evolutionary rescue of a staple monocot crop following a global aphid outbreak provides an opportunity to characterize the genetic and molecular basis (*8*).

Plants adapt to diverse pests via constitutive defenses or induced immunity (*9*). Some defense compounds are natively expressed to deter feeding directly such as glucosinolates, benzoxazinoids, and cyanogenic glucosides (*10–12*). In contrast, induction of common phytohormone and defense networks can result from recognition for various pests (*13*, *14*). *Resistance* (*R*) genes include intracellular receptor proteins which detect molecular patterns and activate defense pathways through initial oxidative and Ca^2+^ bursts, subsequent phytohormone signalling, and ultimately developmental and physiological changes which affect fecundity and/or behavior (*15*, *16*). Intracellular *R* proteins are typically encoded by nucleotide-binding site (NBS) leucine-rich repeat (LRR) receptor genes (NLRs) which directly or indirectly recognize pathogens in the plant cytoplasm (*17*). The NBS and LRR domains of NLRs function to inducibly form oligomeric “resistosomes” (*18–20*). In monocots, NLRs typically have an N-terminal coiled-coil domain (CC), which forms a pentameric calcium channel to activate immunity (*21*). NLRs providing resistance to piercing-sucking herbivores include *Vat*, *Mi-1.2, Adnr1, BPH6, BPH30,* and *BPH40*, however the extent of NLR diversity has not been fully characterized (*22–26*).

Aphids (Hemiptera: Aphididae) feed from the phloem of plants and are one of the fastest colonizing pests (*27*, *28*). Sorghum aphid (*Melanaphis sorghi* Theobald)) (*29*) emerged in 2013 in the Western Hemisphere as a major threat to sorghum [*Sorghum bicolor* (L.) Moench] that quickly spread to most growing regions in the Americas (*30–32*). The *Resistance to Melanaphis sorghi 1* locus (*RMES1*) on chromosome 6 was shown to underlie the evolutionary rescue of global sorghum production and in commercial breeding programs across the Americas after the aphid outbreak (*8*, *33*). *RMES1* is widely deployed for sorghum aphid resistance, however the selection pressure placed on aphid populations highlights the potential for future biotype shifts (*34*). Competing hypotheses on the molecular basis of aphid resistance have been proposed involving cyanogenic toxicity or *R*-gene induced defenses (*8*, *33*, *34*). The cyanogenic glucoside dhurrin is a deterrent of chewing insects and the detoxification gene β-cyanoalanine synthase (*CAS*) is within the *RMES1* sweep region (*8*, *35*, *36*). However, LRR-encoding genes (referred to here on as NLRs) are located within the original mapping interval (*33*). Here we characterize the pangenomic origins and molecular basis of *RMES1*, describing extensive structural variation at a noncanonical NLR gene cluster that led to activated conserved immunity pathways necessary for evolutionary rescue.

## RESULTS

### *RMES1* disrupts aphid feeding to reduce fecundity

The *RMES1* locus was first mapped in a biparental population to Chr06 (*33*) and a separate analysis of a Haitian breeding population under strong aphid infestation identified a selective sweep at *RMES1* (Fig. 1a)(*8*). In order to test mechanistic hypotheses, BC_3_ near-isogenic lines (NIL+, NIL-) were developed with IRAT204 (*RMES1*+ donor) and RTx430 (recurrent parent) which were 94.7% isogenic (Fig. 1a). NIL+ contained a 5.16 Mb introgression from IRAT204 on Chr06 from 2.13 Mb to 7.28 Mb (RTx430v2 reference). The syntenic region on the pangenome reference PI276837 (possessing the resistant *RMES1* SNP S06_02892438) corresponds to 2.06 Mb to 7.25 Mb (PI276837v1) and encompasses the *RMES1* QTL at approximately 2.78–3.21 Mb (PI276837v1, Fig. 1b) (*8*, *33*).

**Fig. 1-.**
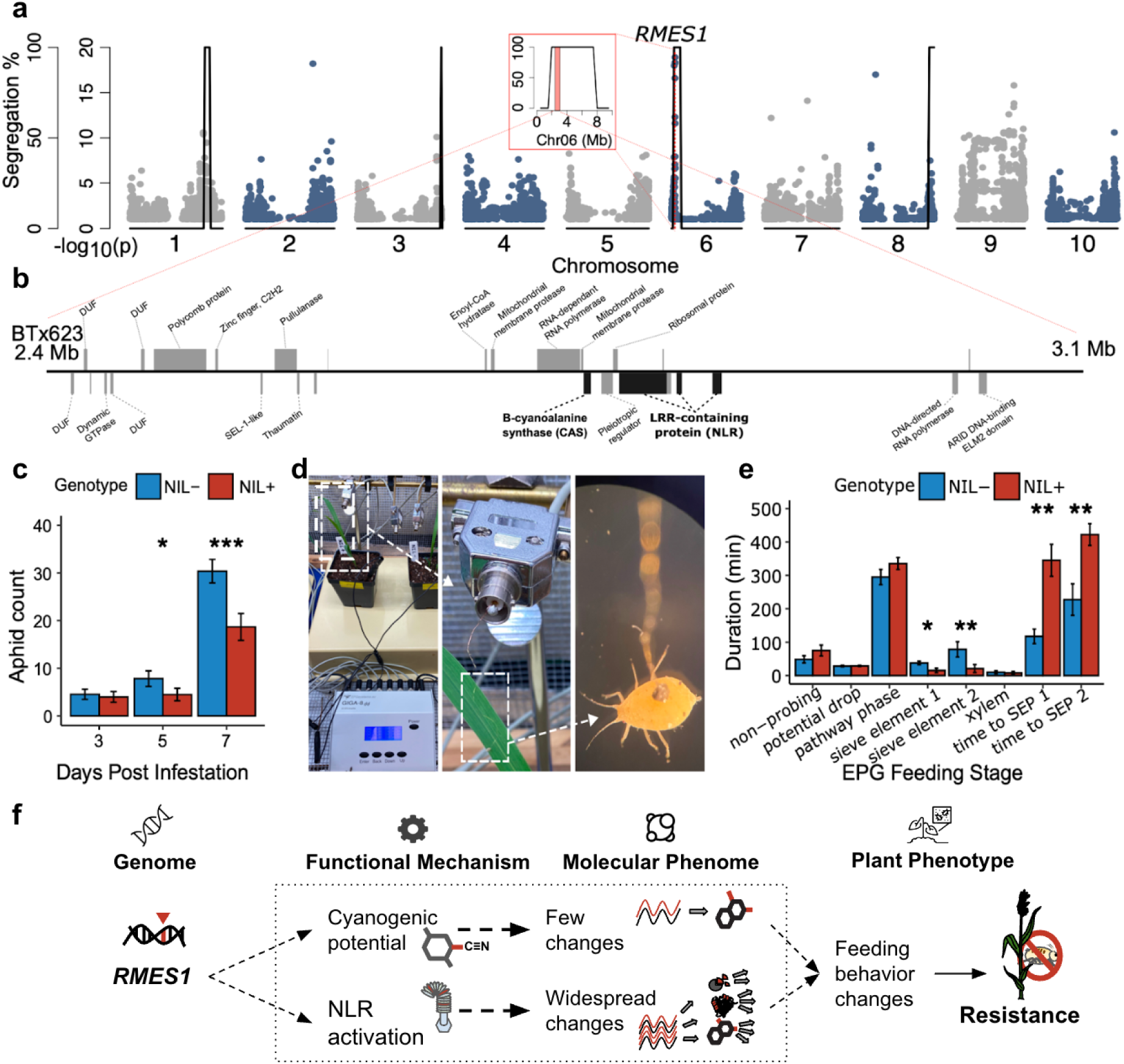
RMES1 reduces aphid fitness by disrupting feeding but the molecular mechanism and pangenomic basis is unknown. a) Selection scan (*F*_ST_) of Haitian breeding population and a global diversity panel (*8*) overlaid with near-isogenic line introgression plot identifies *RMES1* on the RTx430v2 (recurrent parent) sorghum genome. Outer axis for NIL segregation between NIL− (RTx430 recurrent parent haplotype) and NIL+ (donor haplotype). Inner axis for significance (−log_10_(*P*)) of selection scan. Inset highlights Chr06 *RMES1* introgression and QTL from fixation scan as the red shaded region (*8*). b) Genes within *RMES1* QTL (red region of 1a inset) of BTx623v5 highlighting candidate causal genes. Genes above are in forward orientation, below are reverse orientation. Black genes are functional candidates. Unlabeled genes were not assigned automated annotations. c) No-choice assay over 7 day infestation determining aphid fecundity. (* *P*<0.05, ** *P*<0.01, *** *P*<0.001). d) Electrical penetration graph (EPG) experimental setup for detecting feeding behavioral changes. White dotted lines zoom to insect electrode on sorghum leaf and gold wire connected to aphid. e) EPG duration spent at each feeding phase. SEP - sieve element phase. f) Competing functional and molecular hypotheses on the basis of *RMES1* resistance.

To confirm the previously-reported antibiosis resistance trait, a no-choice assay determined aphid fecundity on NILs (Fig. 1c) (*34*). At 5 dpi, *M. sorghi* populations were significantly lower in NIL+ plants compared to NIL− (mean ± se, 4.2 ± 1.3 vs. 7.8 ± 1.6; t-test, *P* < 0.01). At 7 dpi, this effect was stronger (17.1 ± 2.4 vs. 30.4 ± 2.4; *P* < 2e-4). Since NLR-encoding regions from *Medicago truncatula* inhibit feeding of *Acyrthosiphon kondoi* (*37–39*) we next considered if the *RMES1*-mediated fecundity reduction in sorghum aphid is due to altered feeding behavior (Fig. 1d, Data S1). The electrical penetration graph (EPG) technique revealed aphids spent significantly less time in the phloem salivation (E1) and ingestion (E2) phases on NIL+ plants (Fig. 1e). Salivation time (E1) was nearly twice as long on NIL− plants (*P* = 0.02), while phloem ingestion (E2) lasted approximately four times longer on NIL− plants (*P* = 0.02). Additionally, aphids had greater difficulty locating sieve elements on NIL+ plants, taking three times longer to initiate salivation and twice as long to begin ingestion. These results establish that *RMES1* disrupts aphid feeding behavior and reduces fecundity.

### *RMES1* response to infestation causes widespread molecular remodeling

Based on the predicted function of genes at the locus, cyanide detoxification via cyanoalanine synthase and induced immunity via NLRs were determined to be the top candidate mechanisms (Fig. 1b,f). Analysis of the molecular response of uninfested (control), 24 HPI, and 48 HPI NILs found a total of 4,306 detected metabolites and 202 which were significantly affected by genotype or treatment (fold change > 2, *P* value < 0.01). Cyanogenic glucosides, including dhurrin, were not differentially expressed in the metabolome (Data S2). Global expression changes were simultaneously determined using RTx430v2 and 26,120 of 34,601 genes were expressed. Genes regulating the dhurrin pathway were generally downregulated after infestation in NIL+ while being unresponsive NIL− (*35*) (Fig. S1). Notably the *CAS* gene at *RMES1* was not differentially expressed in all comparisons and the biosynthetic steps were downregulated in NIL+. This supports molecular adjustments in response to aphid infestation but not a hydrogen cyanide potential mechanism via *CAS*.

NLR activation typically results in widespread changes to the metabolome and transcriptome which was observed in NIL+ (Fig. 2a) (*40*). Overall, the metabolome principal component 1 (PC1) distinguished infested samples for NIL+ but not NIL− (Fig. 2b). Metabolite functional classes were predicted and diverse phenolic compounds were among those upregulated in NIL+ relative to NIL− (Fig. S2). In response to infestation, hydrolyzable tannins, lignin-related compounds, and flavonoid glycosides were among upregulated metabolites in infested NIL+ (Fig. S3). Fewer metabolites responded in NIL− than NIL+. The diversity of functional classes responsive to infestation suggests multiple metabolic pathways are activated by *RMES1*. Likewise, the transcriptome of infested NILs showed strong evidence of global immunity activation. Over 15% of expressed genes in NIL+ were differentially regulated (N = 8,030, adjusted P < 0.05, fold change > 1.5) compared with 2% in NIL− (N = 610) (Fig. 2c). PC1 captured transcriptional differences in infested NILs to a higher degree in NIL+ (Fig. 2d). PC3 distinguished 24 and 48 HPI timepoints with uninfested samples intermediate (Fig. S4). Differential expression and structure of metabolomes and transcriptomes indicate global induced responses in an *RMES1*-dependent manner.

**Fig. 2-.**
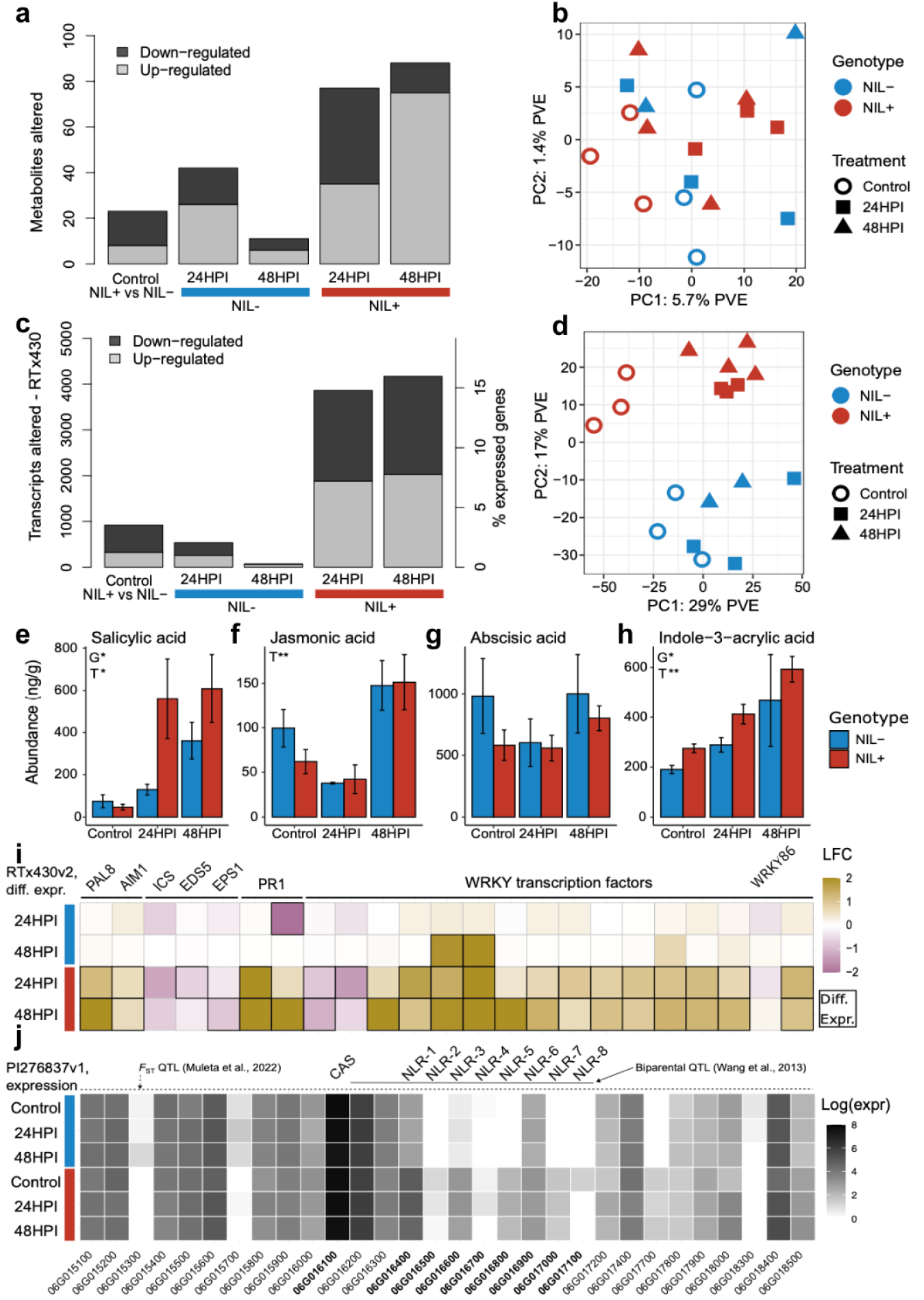
RMES1 broadly remodels the transcriptome and metabolome in response to infestation. a) Differential expression of sorghum metabolomes during aphid infestation identifying *RMES1*-mediated changes (*P* < 0.01, fold change > 2). Control metabolites upregulated in NIL+ relative to NIL-. NIL+/− 24/48 HPI are upregulated in each treatment relative to uninfested controls. b) Principal component analysis (PCA) of metabolomes. PVE - percent variance explained. c) Differential expression of global transcriptome changes during infestation of NILs. Expression was identified using the RTx430v2 recurrent parent for mapping (*P* < 0.05, fold change > 1.5). d) PCA of transcriptomes. e-h) Phytohormone abundance of NILs. Analysis of variance genotype (G) and treatment (T) effects (* *P* < 0.05, ** *P* < 0.01). i) Defense pathway transcriptional induction using RTx430v2 transcriptomes. Salicylic acid marker genes and WRKY transcription factors, outlined boxes indicate differential expression relative to control. j) Log transformed normalized expression of genes at *RMES1* using the PI276837v1 reference. CAS and NLR candidates are labeled above. Note, all gene identifiers are abbreviated from SbPI276837 IDs (e.g. SbPI276837.06G016400 = 06G016400).

We next looked for evidence of defense networks contributing to molecular perturbations and found functional gene enrichment supporting NLR activation. The largest response in co-expression (MEturquoise, N = 2,835) was down regulation after infestation, with NIL+ having a stronger response than NIL− (Fig. S5). This module was enriched for photosynthesis and primary metabolic terms (Data S3). Two modules with continued upregulation over 24 and 48 HPI in NIL+ (MEblue, MEyellow) were enriched for response and signal transduction, cell redox homeostasis, and L-phenylalanine metabolic process terms. Overall, the signalling and metabolic responses are consistent with NLR-based immunity activation and suggest a growth-to-defense transition.

### *RMES1* activates multiple defense signalling pathways

To determine which phytohormones contribute to defense signaling, we quantified salicylic acid, jasmonic acid, 12-oxo-phytodienoic acid (12-OPDA), abscisic acid (ABA), and several auxin conjugates. Salicylic acid, required for NLR-based *Mi-1* aphid resistance, had the strongest response and increased by approximately 10-fold (Fig. 2e-h) (*23*, *41*). Phytohormones more abundant in NIL+ were salicylic acid (ANOVA, *P*<0.05), indole-3-acrylic acid (*P*<0.05), and indole-3-aspartic acid (*P*<0.01) (Fig. S6). Jasmonic acid and its precursor in the jasmonate pathway, 12-OPDA, decreased at 24 HPI in both genotypes before rising indicating a common role in early immunity activation unregulated by *RMES1*. This suggests the causal mechanism at *RMES1* is inducing defense to aphids through salicylic acid signalling.

To confirm the activation of phytohormones signalling by *RMES1*, marker genes were assessed for significant responses (*P*_adj_ < 0.05, fold change > 1.5) (Data S4). For salicylic acid biosynthesis, the PAL pathway was upregulated (*PAL8*, *AIM1*) whereas the ICS pathway was downregulated (*ICS*, *EDS5*, *EPS1*) (Fig. 2i) (*42*, *43*). Signaling involved *PR1* homologs upregulated in a genotype-treatment interaction (Fig. 2i) (*44*). Biosynthesis marker genes for oxylipins including JA, which plays dichotomous roles in sorghum aphid defense signalling (*45*), were differentially regulated in NIL+ (Fig. S7). The related 9-lipoxygenase genes are associated with herbivory defense via death acid biosynthesis (*46*, *47*) and significantly upregulated in NIL+ (*LOX1*, *LOX3*, *LOX4*). WRKY transcription factors play diverse roles in response to environmental stressors and *WRKY86* is the candidate gene underlying *RMES2* aphid resistance (*48*). Despite a strong general response by other WRKY genes, *WRKY86* did not significantly respond to aphid infestation and was slightly downregulated at 24 HPI before upregulating in both genotypes (Fig. 2i, S8). The transcriptional activation of salicylic acid and oxylipin pathways as well as WRKY genes indicate phytohormones and defense networks are activated by *RMES1*.

To investigate candidate genes with improved genome resolution, transcriptomes were mapped to the *RMES1*+ PI276837. Of the eight tandem NLRs in the resistant haplotype, five had negligible expression (< 5 raw reads) in NIL− suggestive of PAV absence (NLR-2, NLR-4, NLR-5, NLR-7, NLR-8) (Fig. 2j). Three NLRs had a significant genotype effect (NLR-1, NLR-3, NLR-5) (Data S4). A putative C2H2-type zinc finger and thaumatin protein (SbPI276837.06G015200, SbPI276837.06G015500) had a significant genotype and interaction effect, however these fell outside of the biparental mapped QTL (*33*). The differential expression of NLRs suggest the strongest candidates for *RMES1* are NLR-1 (SbPI276837.06G016400), NLR-3 (SbPI276837.06G016600) and/or NLR-5 (SbPI276837.06G016800).

### A structural variant at *RMES1* harbors high copy number variation and causal NLRs

Given evidence for an NLR mechanism and differences in gene content at the locus, we next investigated the NLR encoding region across the pangenome. An alignment of BTx623 to PI276837 found a structural variant colocated with the QTL (Fig. 3a). The ∼0.2 Mb insertion coincided with both mapping intervals and five NLRs at the locus, accounting for the difference in NLR copy number between BTx623 and PI276837. To understand the extent of *RMES1* NLR copy number variation (CNV), we searched the pan-proteome for homologous sequences resulting in 163 sorghum orthologs and determined their phylogenetic relationship (Fig. S9). Orthologs were found on Chr06 (*RMES1*) and Chr10. CNV at *RMES1* ranged between two and eight sequences (Fig. 3b). Five clades, established by long well-supported branches, were identified with Chr06 orthologs composed of group-1, group-2, and group-3 (Fig. 3c, Fig. S9). The monophyletic group-1 had low CNV and only two genomes, including PI276837, contained a second paralog. Group-2 and group-3 clades accounted for the majority of NLR CNV (Fig. 3b,c).

**Fig. 3-.**
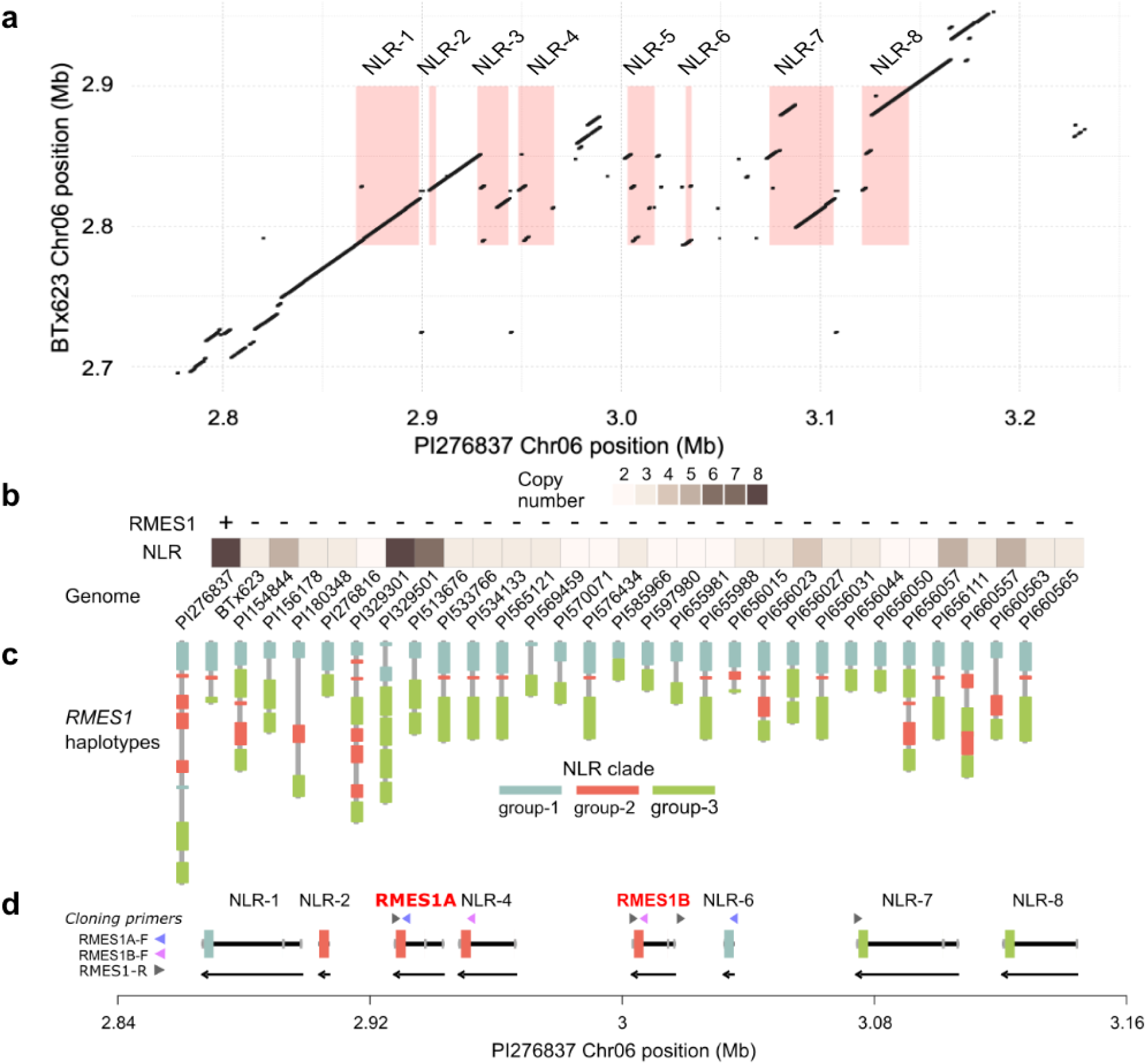
Pangenome structural variant harbors high copy number of NLRs including causal RMES1A and RMES1B. a) Genome alignment of susceptible (BTx623) and resistant (PI276837) haplotypes at *RMES1* encompassing structural variants and NLRs. Red boxes indicate NLR gene locations in PI276837. b) Copy number variation of NLRs at *RMES1* across the sorghum pangenome. c) Pangenome haplotypes of NLR genes at the *RMES1* locus of varying size and genic content. Genes are colored according to phylogenetic clade (Fig. S9) d) PI276837 haplotype array of NLRs with identification of *RMES1A* and *RMES1B* from cloning primers (*50*). Coding sequence is colored by clade, 3’UTR and 5’UTR in grey, intron in black, arrows below indicate reverse strand orientation.

Based on the structural variation coinciding with differential expression (Fig.1, 2), we concluded that NLR-1, NLR-3, and NLR-5 were the likeliest candidates to underlie *RMES1* (*49*). During the preparation of this manuscript, two of these NLRs (NLR-3 and NLR-5; named *RMES1A* and *RMES1B*) were reported in the Chinese sorghum variety Henong 16 as causal for *RMES1* aphid resistance (*50*). Using the cloning primers, NLR-3 (SbPI276837.06G016600) and NLR-5 (SbPI276837.06G016800) were identified as *RMES1A* and *RMES1B*, respectively within the pangenome (Fig. 3d). These genes had been strongly supported by genomic and functional analyses. The Henong 16 haplotype had five tandemly arrayed NLRs and the causal genes were directly adjacent, indicating there are at least two resistant haplotypes.

### *RMES1*-encoded NLR proteins have a noncanonical structure

To better understand the molecular function of the causal NLRs, we investigated their protein structure. Domain and motif prediction identified two LRR regions in all RMES1 paralogs, however, the N-terminus did not appear to encode CC or NBS domains typical of Arabidopsis and monocot NLRs (Fig. 4a). Structural prediction and similarity analysis with AlphaFold and FoldSeek indicated the 298 residues at the N-terminus of RMES1A and RMES1B were highly similar to the NBS domains of canonical NLRs *Arabidopsis thaliana* RPP13L4 (ZAR1, E-value = 1.5e-5) and *Triticum monococcum* CNL9 (Sr35, E-value = 2.8e-5) (Fig. 4b-e, Fig. S10). RMES1A and RMES1B both lack sequence N-terminal to the NBS-like domain, thus they contain neither the typical CC domain, nor a novel domain that could function to activate immunity. The NBS domains of plant NLRs are conserved AAA-ATPase domains which change conformation based on nucleotide binding to a conserved Walker A P-loop motif. The structure of the RMES1 NBS model aligns well with the cryoEM structure of ZAR1, however, the P-loop contains sequence variation (GKT>AAS) in RMES1A and RMES1B that is expected to result in a loss of function (Fig. 4f, Fig. S11) (*51*). Thus, *RMES1* NLRs, despite being causal for aphid resistance, have two unusual features that are expected to independently result in a loss of function for canonical NLRs.

**Fig. 4-.**
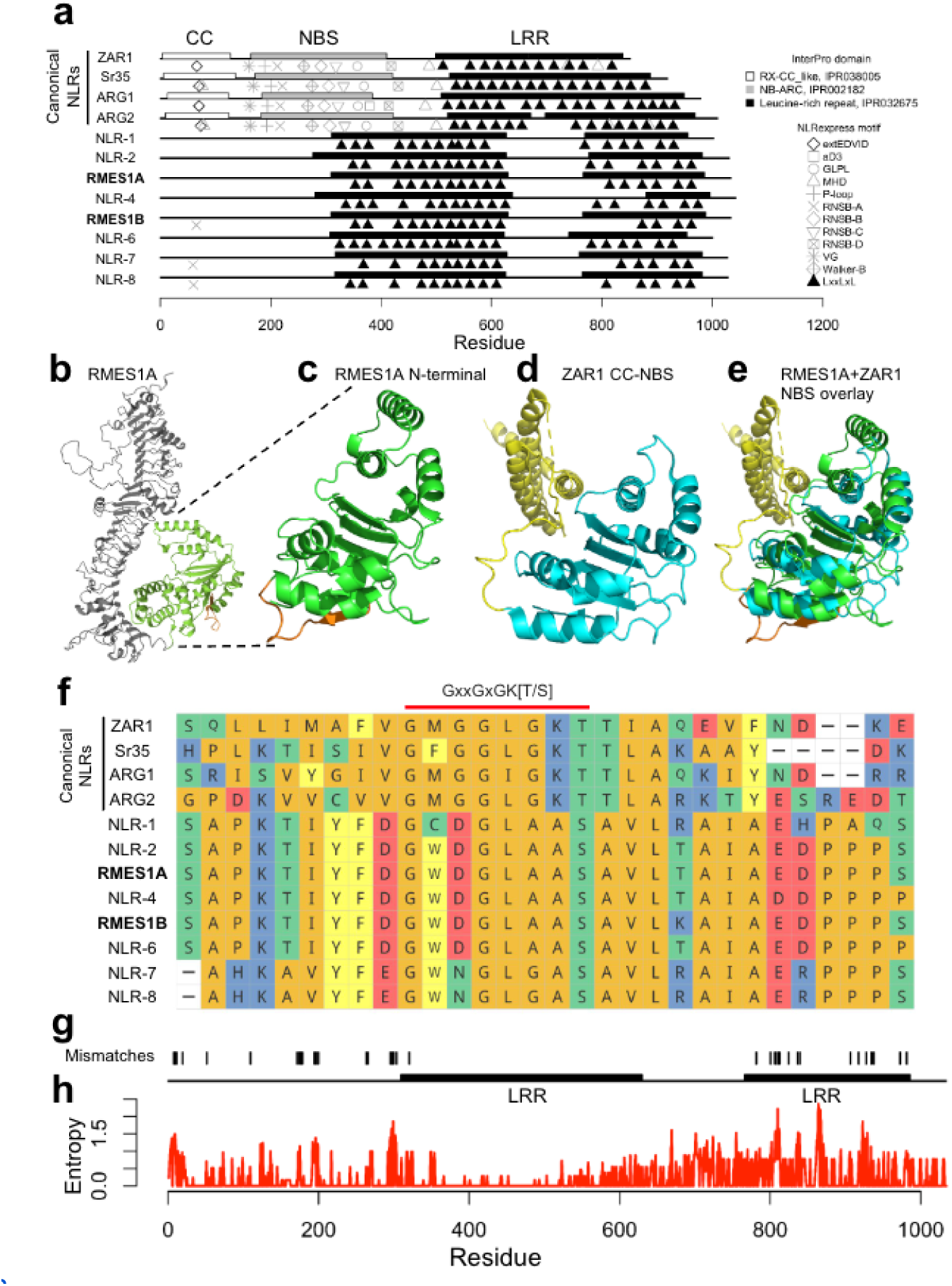
Noncanonical RMES1 NLRs have unusual N-terminal and highly divergent LRR domain. a) Domain and motif annotations of canonical NLRs and noncanonical RMES1 NLRs missing coiled-coil (CC) and NBS domains. b-c) AlphaFold prediction of full length and N-terminal region of RMES1A. LRR region (grey), N-terminal domain (green), and P-loop motif (orange) colored. d) AlphaFold prediction of ZAR1 CC (gold) and NBS domain (blue). e) Close structural alignment of the RMES1A N-terminal domain (green) to the NBS domain of ZAR1 (blue). f) Sequence alignment of canonical NLR P-loop motifs and RMES1 NLRs. g) Positions of residue mismatches between RMES1A and RMES1B. g) Shannon entropy quantifying highly variable residues across a pangenomic allelic series (group-2). Zero indicates invariable residues.

Sequence conservation between the causal proteins and pangenomic orthologs could indicate regions with evolutionary constraints. The PI276837 RMES1A and RMES1B had 96.2% identity (996/1035), however the amino acid residue differences primarily occurred in the NBS-like region or the second LRR domain (Fig. 4g). Phylogenetic clades were used to approximate allelic series and Shannon’s entropy of residues revealed low and highly variable positions (Fig. 4h). Group-1 had fewer variable residues than group-2 and group-3 which both represented highly variable NLRs (>10 positions with entropy bit > 1.5) (Fig. S12). The second LRR domain had notably higher entropy values than the first. The mutated P-loop motif is conserved across the RMES1 pangenome orthologs (entropy = 0). Therefore, it is likely that polymorphisms in the second LRR domain allowed functionalization unrelated to the noncanonical NBS domain.

### Pangenome variation at *RMES1* has ancient evolutionary origin in plants

To trace the origins of *RMES1* beyond the sorghum pangenome, we conducted synteny analysis across the Poaceae super-pangenome. The *RMES1* locus was syntenic to Chr05 of *Brachypodium distachyon*, Chr04 of *Oryza sativa*, and Chr07 of *Setaria viridis* (Fig. 5a). At each syntenic locus, two or more NLRs were adjacent to a β-cyanoalanine synthase CAS ortholog. In order to determine the relationship of RMES1 orthologs across grasses, a phylogenetic tree was built using syntenic and global orthologs from PI276837, BTx623, Setaria, Oryza, and Brachypodium, as well as canonical (*ZAR1*, *Sr35*) and noncanonical NLRs (*BPH6*, *BPH30*, *BPH40*) (*24*, *25*) (Fig. 5b). The physical and phylogenetically adjacent genes of RMES1 support a tandem duplication, while Chr10 orthologs indicate a segmentental duplication originating from Chr06. Interestingly, the brown-planthopper (*Nilaparvata lugens*) resistance gene *BPH40* is a noncanonical NLR allelic to LOC_Os04G08390 and therefore a syntenic ortholog of *RMES1* (*24*). *BPH6* and *BPH30* are more closely related to *RMES1*-like genes than canonical NLRs but are not syntenic to *RMES1*. Poaceae orthologs of *RMES1* share P-loop substitutions (GKT>[A/G]AS) (Fig. S13). In order to determine if *RMES1*-like genes occur outside of Poaceae, a Hidden Markov model developed for the RMES1 NBS-like domain identified 556 orthologs found in Poaceae, Bromeliaceae, and the dicot families of Ranunculaceae, Lauraceae, and Asteraceae, however nearly all (>96%) were found in Poaceae (Fig. 5c).

**Fig. 5-.**
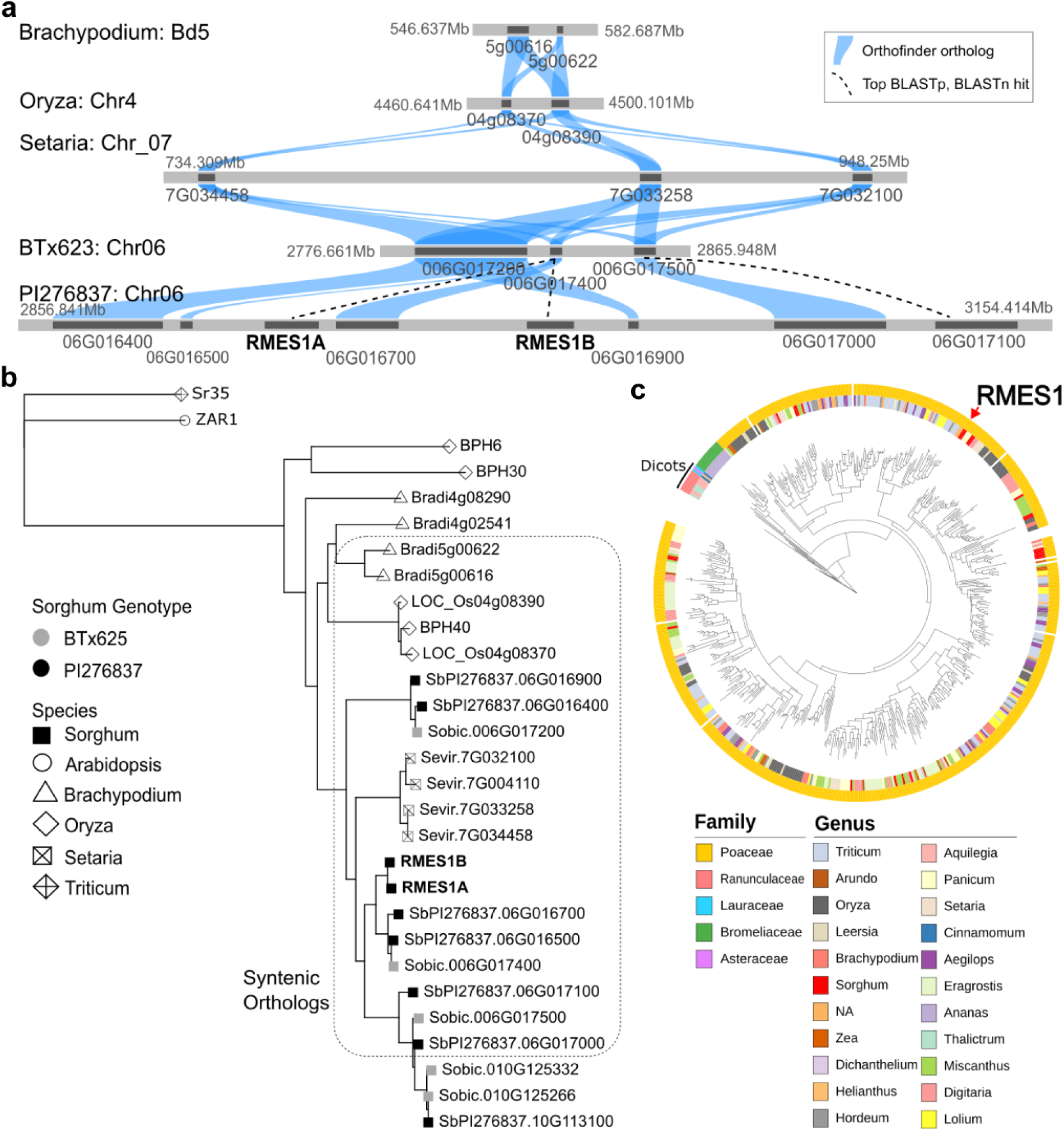
Synteny of RMES1 across the grasses and NLR gene family expansion in plants. a) Syntenic relationships across *B. distachyon*, *O. sativa*, *S. viridis*, and *S. bicolor* (BTx623 and PI276837) genomes trace the ancient origins of the *RMES1*-related NLR gene family in the grass super-pangenome. All genes were within the syntenic orthogroup determined by GeneSpace. Blue connections indicate Orthofinder orthologs, dotted lines are best BLASTp and BLASTn hits for orthogroup members. b) Phylogenetic relationships of RMES1 syntenic and global orthologs, noncanonical BPH NLRs, and canonical NLRs. c) Orthologs of *RMES1* across diverse monocot and dicot plant families identified by an HMM for N-terminal sequence.

We used a k-mer based genotyping approach to determine which genotypes of the pangenome resequencing resource possessed causal RMES1 genes of the PI276837 haplotype. Among 460 landraces across Africa and Asia, only five libraries possessed both RMES1A and RMES1B and were localized to East Africa and Yemen (Fig. 6a). Among all 2,144 resequenced genotypes, 34 libraries possessed both RMES1 genes including ∼10% (13/137) in the resequenced subset of the West African Sorghum Association Panel, which represents breeder’s working collections of local landraces and international breeding lines (Data S5) (*52*). The k-mer analysis conclusive establishes the rarity of the resistant haplotype in global landraces, which was previously hypothesized based on mid-density sequencing data (*8*), as well as establishing the presence of *RMES1* in African breeding networks. The Mendelian nature of this noncanonical NLR activation mechanism coupled with the conserved immunity infrastructure of phytohormone signalling and phytoalexin biosynthesis allowed *RMES1* to be rapidly selected for evolutionary rescue (Fig. 6b,c).

**Fig. 6-.**
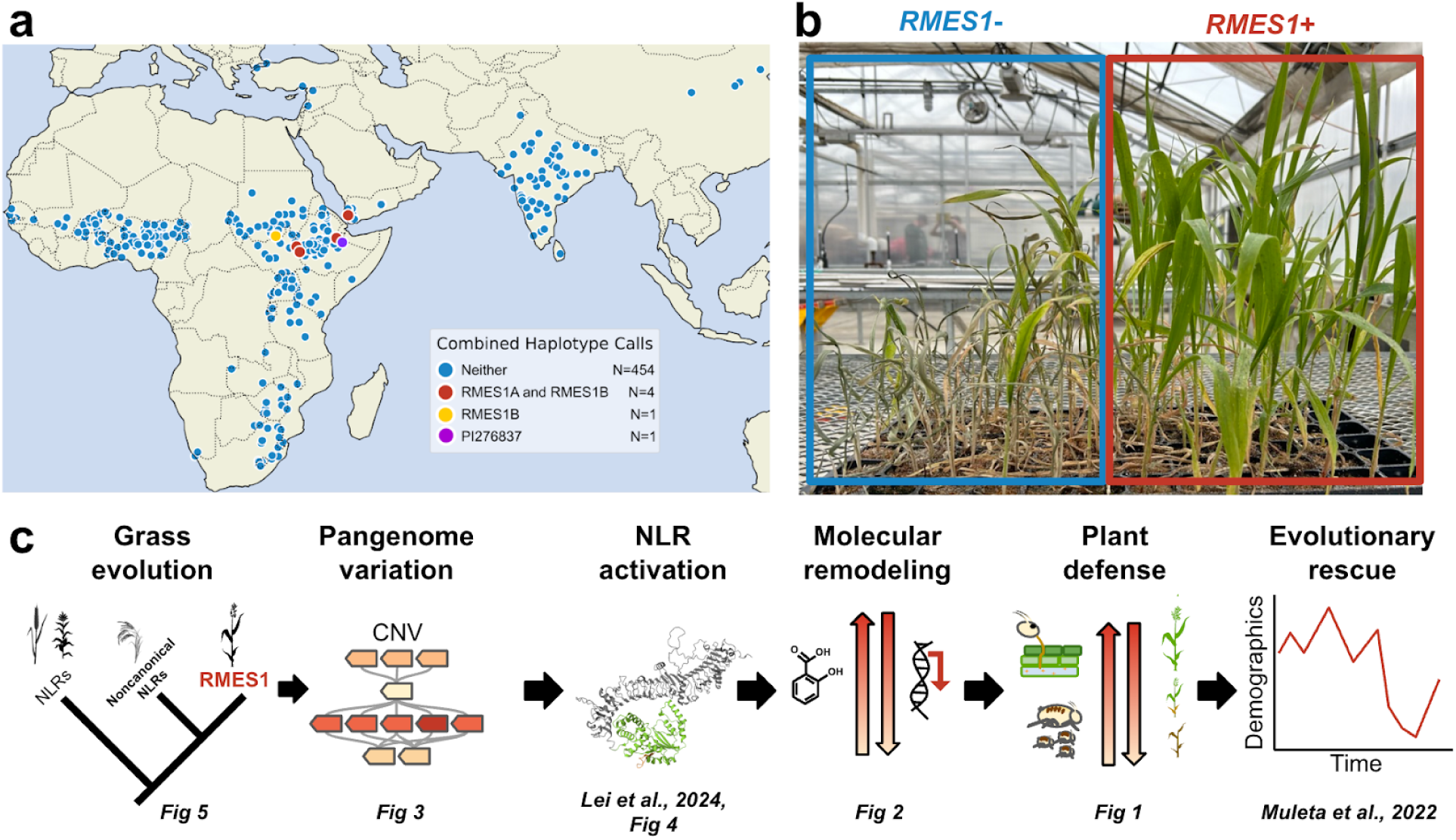
A rare resistant haplotype underlies the genome to phenome map of evolutionary rescue. a) Kmer-based genotyping of *RMES1A* and *RMES1B* in global georeferenced sorghum landrace accessions. b) Near-isogenic lines (NIL− left, NIL+ right) after two weeks of sorghum aphid infestation. c) Integrated genome-phenome model for the evolutionary rescue of sorghum from sorghum aphid by *RMES1*.

## DISCUSSION

### Ancient gene cluster in the super pangenome leading to recent evolutionary rescue

The genomic basis and molecular mechanisms allowing plant populations to rapidly adapt to changing environmental pressures have not been extensively studied. According to Orr’s theory of adaptive walks, a large-effect adaptive variant will be the first to undergo selection when a fitness landscape shifts (*53*). Such variants likely play a larger than expected role in biotic selection where fitness landscapes are more spatially and temporally variable (*54*). NLRs and *R*-genes have been known to play significant roles in local adaptation to pathogenic pressures, however their ability to prevent extirpation through rapid selection have not previously been demonstrated (*55–57*). Here, we show that a large-effect NLR variant activates common immunity networks to disrupt aphid feeding and was responsible for the evolutionary rescue of sorghum.

Early models of adaptive walks only considered *de novo* mutations, however adaptation from standing variation is expected to make a major contribution to regaining fitness (*58*, *59*). Quantitative genetic theory (*60*) and population genomic surveys (*61*) suggest such standing variation is abundant in most populations, and disease resistance QTL often harbors extensive polymorphisms (*62*). Genome organization and expression indicates *RMES1* arose from a locus undergoing gene birth-and-death and highly polymorphic alleles (*63*) (Fig. 3c, 4h, S12). Neofunctionalization often occurs by mutations to the LRR domain leading to novel direct or indirect recognition specificities (*64*, *65*). *RMES1* and its cognate effector directly interact at the LRR domain which has high pangenomic variation (Fig. 4h) (*50*). Notably, *M. sorghi* was first reported on Sudanese sorghum (*29*), suggesting local selection pressure maintained the variant in only a handful of East African landraces, but allowed rapid adaptation to the aphid invasion.

### A noncanonical NLR activates conserved immunity networks

The NLR superfamily is capable of triggering immunity with near limitless inter- and intra-species diversity (*19*). NLR proteins form resistosomes via oligomerization in order to activate their N-terminal signalling domains (*18*). By contrast, RMES1 proteins are unusual in that they lack N-terminal domains and contain variation at the NBS P-loop motif predicted to block oligomerization so must have a non-canonical function distinct from known NLRs (*51*). Artificial mutations to the catalytic lysine are used in molecular investigations of NLR structure and function, however, natural variants for this residue which are functional for immunity have not been observed. RMES1A and RMES1B may form a functional ATPase and/or signalling mechanism as a heterodimer rescuing conformational activation (*66*).

Like *RMES1*, the syntenic brown planthopper resistance gene in *O. sativa* (*BPH40*) appears to require an alternative activation mechanism indicating the noncanonical structure-function relationship is conserved (Fig. S13) (*24*). However, the heterodimer requirement for function appears to be unique to RMES1 as the BPH40 is individually sufficient for function (*50*). The *Vat* cluster in Cucurbitaceae contains several canonical CC-NLRs, however, the only functional resistance identified at *Vat* is in *Cucumis melo* where *Vat-1* and *PmW* alleles provide resistance to melon aphids and potyviruses or powdery mildew, respectively (*67*). *RMES1* and *BPH40* are striking examples of phloem-feeding resistance across Poaceae which may be the conserved ancestral function for this gene family or parallel functionalization in *Sorghum* and *Oryza*.

In support of *RMES1* being a noncanonical NLR with immunity functions, there is strong structural similarity to ZAR1 and Sr35 and molecular outcomes similar to NLR activation in response to aphids (Fig. 2,4) (*41*, *50*, *68*, *69*). Common outcomes of NLR activated immunity include phytohormone signaling leading to phytoalexin production, physiological modification, and potentially hypersensitive response (*70*). For aphid immunity, salicylic acid is required for the NLR *Mi-1.2* resistance in tomato to *M. euphoribae* as well as for *RAG1* resistance to soybean aphid where NLRs are suspected to be causal (*41*, *71*, *72*). Not only did salicylic acid rapidly accumulate in *RMES1* NILs, the PAL pathway appears to be the dominant biosynthetic route and a negative ICS response could indicate it is antagonistic for plant defense (Fig. 2, S6). The 9-lipoxygenase genes controlling death acid production have been shown to play phytoalexin roles in maize to fungal, lepidopteran, and hemipteran pests (*46*, *73*). The activation of these defense mechanisms by *RMES1* demonstrates that, despite its noncanonical structure and unresolved activation mechanism, the downstream functional outcomes were similar to canonical NLRs (Fig. 2) (*50*).

### Using knowledge of pangenome variation to facilitate evolutionary rescue

An emphasis on pangenomic datasets for crops has seen the number of intraspecific *de novo* reference genomes rise over the last decade (*74*). As evolutionary rescue relies on *de novo* or rare variants, it is unlikely that single reference analyses will be sufficient to understand and facilitate such extreme adaptation. Environmental threats, particularly biotic pressures such as insects or pathogens, often require screening extensive diversity in order to identify resistance (*31*, *75*). Pangenomic references will expedite the identification of adaptive traits which are often absent in first-generation single references (Fig. 3a). Further, pangenomic references will allow efficient identification of trait donors even for extremely rare variants as well as movement of alleles between the subpopulations existing for global breeding networks (*76*) (Fig. 6a). As crop intensification and ecological disruption continue, it is important to develop agricultural systems that can withstand the emergence of new stressors (*77*). The proliferation and dispersal of many biotic pests will likely lead to more outbreaks requiring rapid adaptation in host species (*78*). One avenue to addressing food instability is the identification and manipulation of the pangenomic basis and molecular mechanisms that can provide evolutionary rescue. The mechanism of *RMES1* as an immunity activation trigger is a valuable example of how plant populations can rapidly adapt to new threats. However, more genome-phenome elucidation of real-world examples of evolutionarily rescued populations will be needed to learn how pangenome knowledge can facilitate adaptation to global change.

## MATERIALS AND METHODS

### Near-isogenic line development

*RMES1* near-isogenic lines (NILs) were developed with a donor parent IRAT204 (*M. sorghi* resistant, donor) and recurrent backcrossing to RTx430 (*M. sorghi* susceptible). Single plant selections were made at the F_2_ generation of each backcross cycle (BC_x_F_2_) using the KASP marker for *RMES1* (Sbv3.1_06_02892438R, (*8*) and homozygous +/+ plants were backcrossed. *RMES1*-homozygous BC_3_F_4_ lines (NIL+ homozygous for the resistance linked allele, NIL− homozygous susceptible) were used for all experiments described below.

### Aphid behavior and reproduction assays

*M. sorghi* were received from Dr. Scott Armstrong at the USDA-ARS Stillwater, Oklahoma. Aphids were reared on Tx7000 seedlings under laboratory conditions as previously described (*34*). Seedlings were grown in 4.5” pots with soil (Pro-Mix BX) and top layer of greens grade (Profile) to reduce damping off. Colonies were grown in a 46 × 46 × 76 cm cage at 60-70% relative humidity and temperature of 24 ± 1°C (BioQuip Products Inc., Rancho Dominguez, CA). For the no-choice assays, single seedlings were grown in 6” pots. At 3-4 weeks of age or GS1 (*80*), three 4-5 day old apterous *M. sorghi* aphids were placed at the base of the seedlings with a camel hair brush. A clear plastic cylinder was placed over the plant to prevent aphids from leaving the pot with an organdy cloth covering for ventilation. The number of aphids on each plant were counted at the same time of day for a week. For the electrical penetration graph (EPG) experiments to study aphid feeding behavior, individual age-synchronized aphids (4-5 days old) were wired by attaching a thin gold wire (20 µm in diameter) to the dorsum using a conductive silver glue. The wired aphid was gently placed on a four-week-old sorghum plant. Aphid feeding behavior was recorded for eight consecutive hours using an eight-channel Giga-8 DC-EPG amplifier (*81*). The acquired EPG data were annotated using the Stylet+ software. The discoEPG package was employed for calculating feeding parameters and conducting statistical analysis (https://nalamlab.shinyapps.io/test/). The experiment was repeated five times, each including four NIL− and four NIL+ plants. Only data with a total of eight hours of recording were kept for further analysis. A total of 14 NIL− and 13 NIL+ plants were used in the analysis.

### Infestation timecourse sampling for transcriptomic and metabolomic analysis

A 2 × 3 factorial design was used with NIL+ and NIL− plants infested for 24 (24 hours post infestation, 24 HPI) and 48 hours (48 HPI) as well as an uninfested (control) sample collected at the same time as the 48 HPI sample. A 50 ml falcon tube with the conical end cut off and a hole with organdy cloth on the cap was placed over the third true leaf of two plants and infested with twenty adult apterous aphids. Cotton balls were used to cover the open end of the tube to ensure maximum response from aphid feeding. Control samples were handled the same way but were not infested. Samples were collected at 12 PM after 24 and 48 hours by pooling both leaves. The bottom (basal) inch of tissue from both leaves were flash frozen for sequencing. The remainder of the sample was flash frozen for HPLC analysis. There were 4 replicates collected, however, RNA sequencing analysis identified genetic contamination and six samples were removed from all analyses leaving NIL+ control (N = 3), NIL+ 24 HPI (N = 3), NIL+ 48 HPI (N = 4), NIL− control (N = 3), NIL− 24 HPI (N = 3) and NIL− 48 HPI (N = 2) (Fig. S14).

### RNA sequencing and analysis

RNA was extracted using Zymo Quick-RNA minipreps (Thermo Scientific) and treated with DNAse using Invitrogen Turbo DNA-free kit (Thermo Scientific). RNA quality was checked using a NanoDrop 2000 (Thermo Scientific) and ∼2 μg was submitted on dry-ice to Novogene Corporation Inc. (2921 Stockton Blvd, Sacramento, CA, US, 95817), where RNA integrity was confirmed using an Agilent 2100 Bioanalyzer (RIN > 7). Samples were sequenced on an Illumina NovaSeq 6000 Sequencing System with 150 bp paired-end reads with ∼54.7 M reads per sample. Trimmed and quality filtered reads generated by Novogene were aligned to the PI276837 (https://phytozome-next.jgi.doe.gov/info/SbicolorPI_276837_v1_1) and RTx430v2 (https://phytozome-next.jgi.doe.gov/info/SbicolorRTx430_v2_1) reference genomes for transcriptome analysis and variant calling (*82*). Reference genomes were indexed and aligned to with STAR v2.7.10 using 2-pass mode (*83*). For variant calling, BAM files were processed using the following in GATK v 4.2.5.0 unless otherwise noted: Duplicates were marked and read groups were added using Picard, reads were split using SplitNCigarReads, variants were called individually using HaplotypeCaller, gVCFs were combined using CombineGVCFs, and jointcalled VCF files were produced with GenotypeGVCFs (*84*). Finally, VCFs were filtered using VariantFiltration (QUAL > 30, SQR > 3, FS > 60, MQ < 40). VCFs were analyzed in base R v4.4.1 with variants summarized over 0.5 Mb windows and segregating markers were used to estimate segregation percentage between NILs.

For transcriptome analysis the transcript abundance was quantified using FeatureCounts v2.0.1 (*85*). Differential gene expression was determined using DESeq2 v1.38.1 in R v4.2.2 (*86*, *87*). After removal of contaminated samples, dispersion estimate results suggest the DESeq2 model assumptions were satisfied (Fig. S15). Principal component analysis was performed with R/prcomp and plotted with ggplot2 v3.4.1 (*88*). Coexpression analysis was done using Weighted Gene Coexpression Network Analysis (WGCNA) with the expressed transcriptome (*89*). Coexpression modules were visualized using GGally in ggplot2 (*88*). Genes were determined to be significantly differentially expressed with a Benjamini-Hochberg adjusted *p*-value < 0.05 and fold change greater than 1.5 (L2FC > 0.58). Analysis of potential causal gene and pathway candidates were determined *a priori* through literature search and SorghumBase orthology finder (Data S4)(*90*). Genes involved in aphid defense and/or phytohormone signaling in Arabidopsis or other species were searched on SorghumBase and all orthologous genes in BTx623v5 were considered as candidate genes (Data S1). Lipoxygenase and jasmonate ZIM-domain gene families were characterized in sorghum previously and Sobic IDs were already available (*91*, *92*). Global, phytohormone, and defense transcriptome pathways were analyzed using RTx430v2. Transcriptome and genomic analysis of the *RMES1* region was done with PI276837. For analysis of potential causal genes, all 35 PI276837 genes between 2.5–3.5 Mb were considered.

### Phytohormone and untargeted metabolome analysis

Phytohormone and metabolome analysis was conducted (on the same samples as transcriptome analysis) following (*93*) with full modifications described in the Supplemental Methods. Briefly, frozen samples were lyophilized, homogenized, and extracted from 30 mg of tissue with 80% methanol. The extracts were analyzed with UPLC-MS/MS for general phytohormone profiling and untargeted analysis. Authentic standards used in this assay included jasmonic acid-d5 (JA-D5), jasmonic acid (JA), salicylic acid-D4 (SA-D4), salicylic acid (SA), indole-3-acrylic acid (IACA), indole-3-carboxylic acid (ICA), indole-2,4,5,6,7-d5-3-acetic acid (IAA-D5), 12-oxo-phytodienoic acid (OPDA), IA-aspartic acid (IA-Asp), abscisic acid-D6 (ABA-D6), L-alanine-^13^C_3_, L-phenylalanine-^13^C_6_, fumaric acid-^13^C_4_, L-tryptophan-^13^C_11_, and indole-3-acetic-acid-^13^C_6_.

Phytohormones that were detected and quantified include SA, JA, ABA, OPDA, IAA, ICA, IACA, and IA-Asp. UPLC-MS/MS analysis was performed on a Waters ACQUITY Classic UPLC Premier T3 1.7μm coupled to a Waters Xevo TQ-S triple quadrupole mass spectrometer. Raw data was analyzed with Skyline v21 open source software for retention time and peak area integration (*94*). Peak areas were extracted for target compounds detected in biological samples and normalized to the peak area of the appropriate internal standard or surrogate in each sample. Absolute quantitation (ng/g) was calculated using the linear regression equation generated for each compound from the calibration curve.

Untargeted metabolomes were quantified on a Waters Acquity UPLC system. Separation was achieved using a Waters ACQUITY UPLC Premier T3 1.7μm Column, using a gradient from solvent A (0.1%formic acid in water) to solvent B (0.1% formic acid in acetonitrile) and a flow rate of 0.5 mL/min. The column eluent was infused into a Waters Xevo G2-XS Q-TOF-MS with an electrospray source in negative ionization sensitivity mode, with MS^E^ data independent MS/MS acquisition. XCMS (*95*, *96*) version 3.20.0 was used to process raw data using R v4.2.2. RAMClustR version 1.2.4 in R version 4.2.2 was used to normalize, filter, and group features into spectra (*97*). MSFinder (*98*) was used for spectral matching, formula inference, and tentative structure assignment. Results were imported into the RAMClustR object. A total score was calculated based on the product scores from the findmain function and the MSfinder formula and structure scores. A total of 14,130 annotation hypotheses were tested for 4306 compounds. The highest total score was selected for each compound, considering all hypotheses. Direct parent class was used in the manuscript, full annotation is reported in Supp. Data 2. For statistical analysis of phytohormones, a two-way ANOVA was performed, fitting a linear model with genotype, timepoint, and their interaction. Pairwise statistical comparisons of metabolite abundance was conducted using Welch’s *t*-test (two-sided, unequal variance) in R. Fold change (FC) was calculated as the ratio of group means, and Benjamini–Hochberg adjusted *p*-values were reported.

### Comparative genomics analyses

Intraspecific orthology of *RMES1-*like NLRs was determined using the sorghum pan-proteome (https://phytozome-next.jgi.doe.gov/sorghumpan/). Sequences were clustered using MMseqs2 (*99*) using the following parameters: minimal sequence identity 50%, coverage 80%, coverage mode 1 (--min-seq-id 0.5 -c 0.8 --cov-mode 1). All *RMES1* NLR candidates were found in a single cluster. Cluster sequences were aligned using mafft v7.505 (*100*) with default parameters and cluster phylogeny was constructed using RAxML-NG (*101*) with the following options: model JTT, 100 bootstrap replicates (raxml-ng --all --bs-trees 100 --model JTT). The resulting tree had five well-supported (bootstrap >95%) clades corresponding to group-1, group-2, and group-3 sequences on Chr06 and group-4 and group-5 on Chr10. We extracted the corresponding sub-alignments and calculated per-position Shannon entropy as previously described (*102*).

Orthology of sorghum genes across Poaceae was determined using GENESPACE with default specifications (*103*). Briefly, GENESPACE applies OrthoFinder (*104*) to the protein sequences of 30 sorghum pangenome members, *Setaria*, *Brachypodium*, and *Oryza*, then parses the blast hits to syntenic regions. The resulting syntenic and phylogenetically hierarchically orthologous sets of genes (‘orthogroups’, ‘OGs’) were used for phylogenetic comparisons. Copy number variation was identified as genes belonging to the same OG and shared in both BTx623v5 and PI276837 but at different copy numbers (*105*). Syntenic orthologs from GENESPACE were then used in ‘riparian’ plots to track their positions across genomes. To determine gene locations within repeat expansions, we mapped 100 bp windows with 50 bp overlaps from query to reference target sequences using DEEPSPACE (github.com/jtlovell/DEEPSPACE), which calls minimap2 on windowed sequences. The resulting dotplots can be integrated with positional coordinates of genes to understand broader patterns of sequence expansion. The presence of *RMES1*-like genes outside of *Poaceae* was determined with an HMM of the noncanonical N-terminal. The original cluster alignment was reduced manually using jalview to remove gappy columns and gappy sequences and was trimmed to correspond to the N-terminal NB-ARC-like domain and an HMM was produced. This was used to search the UniRef90 database with SeqKit (*106*).

### Protein motif/domain analyses and structural modeling

Protein sequences for PI276837 RMES1 NLRs, Sobic.007G085400 (*ARG1*), and Sobic.005G047700 (*ARG2*) as well as canonical AtZAR1 (AEE78731.1) and TaSr35 (AGP75918.1) were used to determine motif and domain presence with NLRexpress and InterPro InterPro v101.0 (*107*, *108*). We used AlphaFold2 (*109*) as implemented in ColabFold (*110*) to predict the structures of NLR candidates at *RMES1*. Structural homologs of the N-terminal non-LRR domain of *RMES1* NLR candidates were identified using Foldseek (*68*). Structures were visualized in Chimera (*111*).

### K-mer genotyping

K-mer genotyping of the *RMES1* locus from *Sorghum bicolor* reference PI276837 was conducted within Illumina resequencing libraries (N = 2,143). Single copy 80bp k-mers (80mers) were extracted from genes SbPI276837.06G016600 and SbPI276837.06G016800, generating 15,507 and 13,507 80mers, respectively. To ensure these 80mers were (i) unique to PI276837 and (ii) could be genotyped in short-read Illumina data (i.e. ‘typable’ markers), single copy 80mers were searched using exact matches in Illumina polishing libraries for each pan-genome reference. Only 80mers present in PI276837 and absent in all other references with a minimum observed count of one, and a maximum observed count of 100, were retained. This search yielded 3,586 typable 80mers, 1,322 for SbPI276837.06G016600 and 2,264 for SbPI276837.06G016800. Next, the typeable markers were used to genotype all re-sequenced individuals using the same exact match criteria as the Illumina reference polishing libraries. Based on the histogram distribution of the number of kmer matches per gene, genes with more than 95% of total typable matches were considered present within that library.

## ACKNOWLEDGEMENTS

We thank Joint Genome Institute and John Vogel for the use of the RTx430v2 reference genome. This study is made possible by the support of the American People provided to the Feed the Future Innovation Lab for Collaborative Research on Sorghum and Millet through the United States Agency for International Development (USAID) under associate award no. AID-OAALA-16-00003, “Feed the Future Innovation Lab for Genomics-Assisted Sorghum Breeding”. The contents are the sole responsibility of the authors and do not necessarily reflect the views of USAID or the United States Government. CJV’s contribution was supported by the Samuel and Hazel Litzenberger Fellowship and by the Bill and Melinda Gates Foundation through the grant “Green Evolution - Accelerating Dryland Cereals Improvement for Africa (INV-053669)” to GPM. Genomic data generation was supported by The Bill and Melinda Gates Foundation through their grant “Sorghum Genomics Toolbox: a TERRA collaboration (INV-009319)” to Todd Mockler (Donald Danforth Plant Science Center).

