## Supplementary Materials for "Ancient pangenomic origins of noncanonical NLR genes underlying the recent evolutionary rescue of a staple crop"

### Phytohormone analysis

**Sample Preparation**

The authentic standards used in this assay included jasmonic acid-d5 (JA-D5), jasmonic acid (JA), salicylic acid-D4 (SA-D4), salicylic acid (SA), indole-3-acrylic acid (IACA), indole-3-carboxylic acid (ICA) were purchased from Sigma-Aldrich (St. Louis, MO, USA). Indole-2,4,5,6,7-d5-3- acetic acid (IAA-D5) were purchased from CDN Isotopes (Canada). 12-oxo-phytodienoic acid (OPDA) was purchased from Cayman (Ann Arbor, MI). IA-aspartic acid (IA-Asp) was purchased from Toronto Research Chemicals (Canada). Abscisic acid-D6 (ABA-D6) was purchased from Olchemim (Czech Republic). L-alanine13C3, L-phenylalanine13C6, fumaric acid13C4, Ltryptophan13C11, and indole-3-acetic-acid13C6 were purchased from Cambridge Isotope Laboratories (MA, USA).

The phytohormone analysis was conducted following reference [1] with modifications described below. Frozen samples were lyophilized. The dried samples were added with stainless steel balls and homogenized in the Bullet Blender for 2 min. The homogenate (about 30 mg) was weighed into Eppendorf tubes and added with 1 mL of cold 80% methanol in water, 20 µL of phytohormone internal standards (500 ng/mL of SA-d4, 1000 ng/mL of ABA-d6, 500 ng/mL of JA-d5, 200 ng/mL of IAA-d5 in 50% methanol), and 10 µL of internal standards for untargeted analysis (600 µg/mL L-alanine13C3, 60 µg/mL L-phenylalanine13C6, 30 µg/mL fumaric acid13C4, 300 µg/mL Ltryptophan13C11, and 30 µg/mL indole-3-acetic-acid13C6 in 50% methanol). The mixture was vigorously mixed for 0.5 h, followed by 15 min of sonication, and another 0.5 h of mixing. Then the mixture was centrifuged at 15,000 g and 4°C for 15 min. Supernatants were recovered, of which 0.1 mL was saved for untargeted analysis and 0.8 mL was transferred to a new vial. To the remaining pellets, 0.5 mL of 80% methanol was added, and the sample was mixed for 15 min. The supernatant after centrifugation was combined with first aliquot of 0.8 mL, dried down, and resuspended in 100 µL of 50% methanol for phytohormone analysis. A small aliquot (20 µL) was taken from each study sample and pooled to generate a quality control (QC) sample. Sample extracts and QCs were stored at -80°C until analysis.

**UPLC-MS/MS Analysis**

UPLC-MS/MS analysis was performed on a Waters ACQUITY Classic UPLC coupled to a Waters Xevo TQ-S triple quadrupole mass spectrometer. Chromatographic separations were carried out on a Waters ACQUITY HSS T3 column (2 x 50 mm, 1.7 µM). Mobile phases were (A) water with 0.1% formic acid and (B) acetonitrile with 0.1% formic acid. The LC gradient was as follows: time = 0 min, 1% B; time = 0.65 min, 1% B; time = 2.85 min, 99% B; time = 3.5 min, 99% B; time= 3.55 min, 1% B; time =5 min, 1% B. Flow rate was 0.5 mL/min and injection volume was 3 µL. Samples were held at 6°C in the autosampler, and the column was operated at 45°C. Mass detector was operated in ESI+ and ESI- mode. The capillary voltage set to 0.7 kV. Inter-channel delay was set to 3 msec. Source temperature was 150°C and desolvation gas (nitrogen) temperature 450°C. Desolvation gas flow was 1000 L/h, cone gas flow was 150 L/h, and collision gas (argon) flow was 0.15 mL/min. Nebulizer pressure (nitrogen) was set to 7 Bar. The MS acquisition functions were scheduled by retention times. Autodwell feature was set for each function and dwell time was calculated by Masslynx software (Waters) to achieve 12 points-across-peak as the minimum data points per peak. The retention time, MRM transitions, cone and collision energy of each compound were described in spreadsheet “transitions”.

Data Processing

All Raw data files were imported into the Skyline open source software package [2]. Each target analyte was visually inspected for retention time and peak area integration. Peak areas were extracted for target compounds detected in biological samples and normalized to the peak area of the appropriate internal standard or surrogate in each sample. Absolute quantitation (ng/g) was calculated using the linear regression equation generated for each compound from the calibration curve.

Calibration Curves

Reference standard at various concentrations was mixed with internal standard in methanol for calibration curve. Normalized peak areas are plotted against expected concentrations. See “curve” in the result sheet for details.

Quality Control

Study specific QC samples were generated representing a pool of all samples. The QC samples was injected after every 6 samples, and the coefficient of variance (CV) of QCs was used to evaluate instrument stability. The CV of all detected compounds were <11% (n=5). See attached spreadsheet “QC” for details.

#### Untargeted Metabolome analysis

**Sample Preparation:**

The authentic standards used in this assay included jasmonic acid-d5 (JA-D5), jasmonic acid (JA), salicylic acid-D4 (SA-D4), salicylic acid (SA), indole-3-acrylic acid (IACA), indole-3-carboxylic acid (ICA) were purchased from Sigma-Aldrich (St. Louis, MO, USA). Indole-2,4,5,6,7-d5-3-acetic acid (IAA-D5) were purchased from CDN Isotopes (Canada). 12-oxo-phytodienoic acid (OPDA) was purchased from Cayman (Ann Arbor, MI). IA-aspartic acid (IA-Asp) was purchased from Toronto Research Chemicals (Canada). Abscisic acid-D6 (ABA-D6) was purchased from Olchemim (Czech Republic). L-alanine-13C3, L-phenylalanine-13C6, fumaric acid-13C4, L-tryptophan-13C11, and indole-3-acetic-acid-13C6 were purchased from Cambridge Isotope Laboratories (MA, USA).

Pooled Quality Control Sample preparation:

A small aliquot (20 µL) was taken from each study sample and pooled to generate a quality control (QC) sample. Sample extracts and QCs were stored at -80°C until analysis.

Blank sample preparation:

A blank sample was taken through the entire sample extraction process to enable recognition and removal of contaminant and system peaks.

**UPLC-MS Analysis:**

Sample run order was fully randomized, with a pooled QC sample injected approximately every 7 injections. The process (extraction) blank was also injected last. One microliter of each sample was injected onto a Waters Acquity UPLC system. Separation was achieved using a Waters ACQUITY UPLC Premier T3 1.7µm Column (2.1 x 100 mm), using a gradient from solvent A (0.1% formic acid in water) to solvent B (0.1% formic acid in acetonitrile) and a flow rate of 0.5 mL/min. The 20 minute long separation gradient used is shown in the table below:

| Time (minutes) | Buffer B% |
|---|---|
| 0–1 | Hold, 1% |
| 1—13 | Linear Gradient, 1% to 98% |
| 13—16 | Hold, 98% |
| 16—16.05 | Linear Gradient, 98% to 1% |
| 16.05—20 (end) | Hold, 1% |

The column and samples were held at 45 °C and 6 °C, respectively. The column eluent was infused into a Waters Xevo G2-XS Q-TOF-MS with an electrospray source in negative ionization sensitivity mode, with MSE data independent MS/MS acquisition. The following parameters were used for MS1 scan: 50-1200 m/z mass range with 0.1 seconds per scan, collision energy 6 V. MSE acquisition occurred at a scan rate of 0.1 seconds, mass range

50-1200 m/z, and collision energy was ramped from 15 to 30 V. Calibration was performed using sodium formate with 1 ppm mass accuracy. The capillary voltage was held at 700 V in positive mode or 2200 V in negative mode. The source temperature was held at 150 °C and the nitrogen desolvation temperature at 450 °C with a flow rate of 1000 L/hr. Lockspray reference mass was used to correct for drift, with 40 seconds interval between scans, 0.1 seconds/scan and signal averaged over 3 scans. LeuEnk was used for mass correction, with reference masses of either positive 556.2771 m/z or negative 554.2615 m/z.

Data Analysis and Statistics:

XCMS (Smith 2006, Tautenhahn 2008, v3.16.1) version 3.20.0 was used to process raw data using R v4.2.2. The following processing steps were used: (1) Peak detection (CentWave) : ppm = 30 peakwidth = c(2.5, 15), snthresh = 15, prefilter = c(3, 50), mzCenterFun = wMean, integrate = 1, mzdiff = 0.01, fitgauss = TRUE, noise = 2, verboseColumns = TRUE, roiList = list(), firstBaselineCheck = TRUE, roiScales = numeric(0), extendLengthMSW = TRUE. (2) Peak grouping (PeakDensity) : bw = 2.5, minFraction = 0.7, minSamples = 1, binSize = 0.015, maxFeatures = 50. (3) Retention time correction (PeakGroups) : minFraction = 0.5 extraPeaks = 1, smooth = loess, span = 0.2, family = gaussian, subset = integer(0), subsetAdjust = average. (4) Peak grouping (PeakDensity) : bw = 1.5, minFraction = 0.75, minSamples = 1, binSize = 0.015, maxFeatures = 50. (5) Missing peak filling (FillChromPeaks) : expandMz = 0 expandRt = 0, ppm = 0, fixedMz = 0, fixedRt = 0.

RAMClustR version 1.2.4 in R version 4.2.2 was used to normalize, filter, and group features into spectra.XCMS (Smith 2006)(Tautenhahn 2008) output data was transferred to a ramclustR object using the rc.get.xcms.data function. Feature data was extracted using the xcms featureValues function. Features with missing values were replaced with small values simulating noise. For each feature, the minimum detected value was multiplied by 0.5. Noise was then added using a factor of 0.5. The absolute value of this value was used as the filled value to ensure that only non-negative values carried forward. Variance in quality control samples was described using the rc.qc function within ramclustR. Summary statistics are provided including the relative standard deviation of QC samples to all samples in PCA space, as well as the relative standard deviation of each feature/compound in QC samples, plotted as a histogram. Features were normalized by linearly regressing run order versus qc feature intensities to account for instrument signal intensity drift. Only features with a regression p-value less than 0.05 and an r-squared greater than 0.1 were corrected. Of 44758 features, 4600 were corrected for run order effects. Features which failed to demonstrate signal intensity of at least 2-fold greater in QC samples than in blanks were removed from the feature dataset. 11951 of 44758 features were removed. Features were filtered based on their qc sample CV values. Only features with CV values less than or equal to 0.5 in MS or MSMSdata sets were retained. 5426 of 32807 features were removed.

Features were clustered using the ramclustR algorithm (Broeckling 2014). Parameter settings were as follows: st = 2.74, sr = 0.5, maxt = 274, deepSplit = FALSE, hmax = 0.3, minModuleSize = 2, and cor.method=pearson. Molecular weight was inferred from in-source spectra (Broeckling2016) using the do.findmain function, which calls the interpretMSSpectrum package (Jaeger 2016). Parameters for do.findmain were set to: mode = positive, mzabs.error = 0.005, ppm.error = 10, ads = [M+H]+ [M+Na]+ [M+K]+ [M+NH4]+ [2M+H]+ [2M+Na]+ [2M+K]+ [2M+NH4]+ [3M+H]+ [3M+Na]+ [3M+K]+ [3M+NH4]+, nls = [M+H-COCH2]+ [M+H-C2H3NO]+

[M+H-H2O]+ [M+H-NH3]+ [M+H-HCOOH]- [M+H-C6H12O6]+ [M+H-C5H10O5]+ [M+H-C12H22O11]+.

MSFinder (Tsugawa 2016) was used for spectral matching, formula inference, and tentative structure assignment. Results were imported into the RAMClustR object. A total score was calculated based on the product scores from the findmain function and the MSfinder formula and structure scores. A total of 14130 annotation hypotheses were tested for 4306 compounds. A complete spreadsheet of all annotation hypothesis and scores can be found in the 'spectra/all.annotations.csv' file, and a subset of only those selected for annotation can be found in the 'spectra/assigned.annotations.csv' file. Spectra matches took precedence over computational inference based annotations. The following database(s) were assigned as 'priority': chebi, coconut. The database priority.factor was set to 0.9 to decrease scores for compounds which failed to match priority database(s). The list of 8396 inchikeys is provided. The inchikey priority.factor was set to 0.9 to decrease scores for compounds with non-matching inchikey(s). The highest total score was selected for each compound, considering all hypotheses.

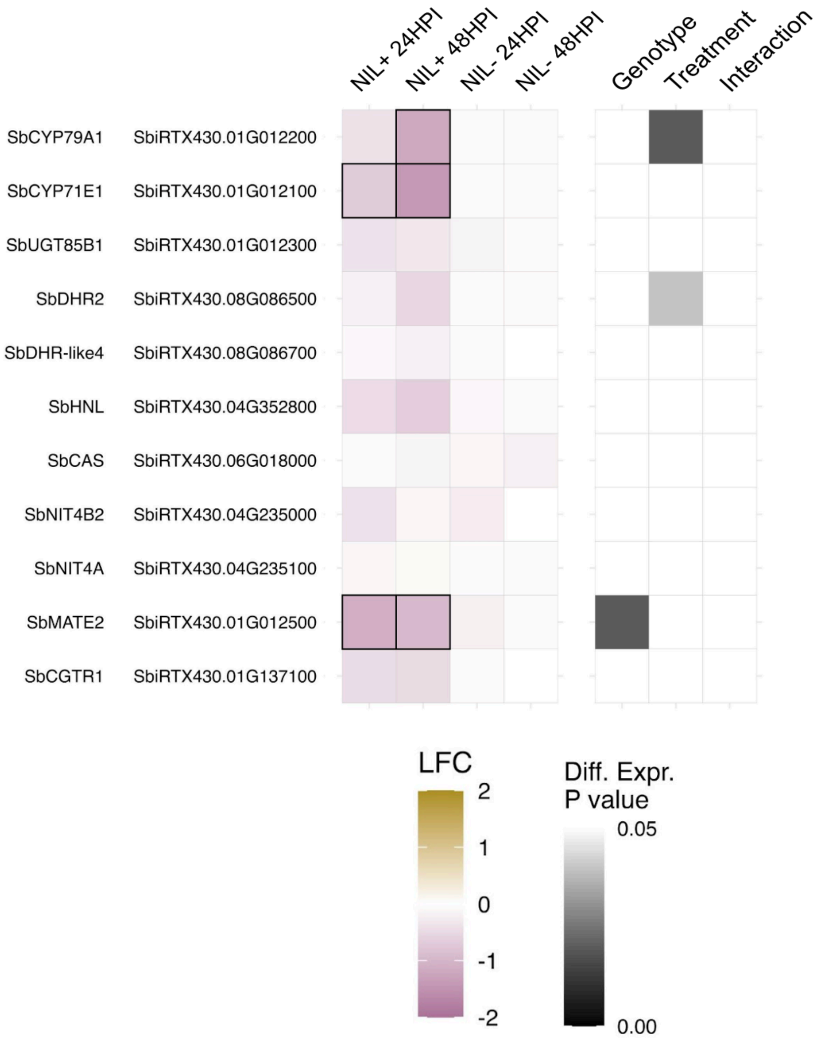


**Figure S1: Differential expression of sorghum genes (reference RTx430) in the dhurrin pathway in response to infestation.** Log fold change (LFC) is in relation to uninfested controls of the same genotype. HPI, hours post infestation.

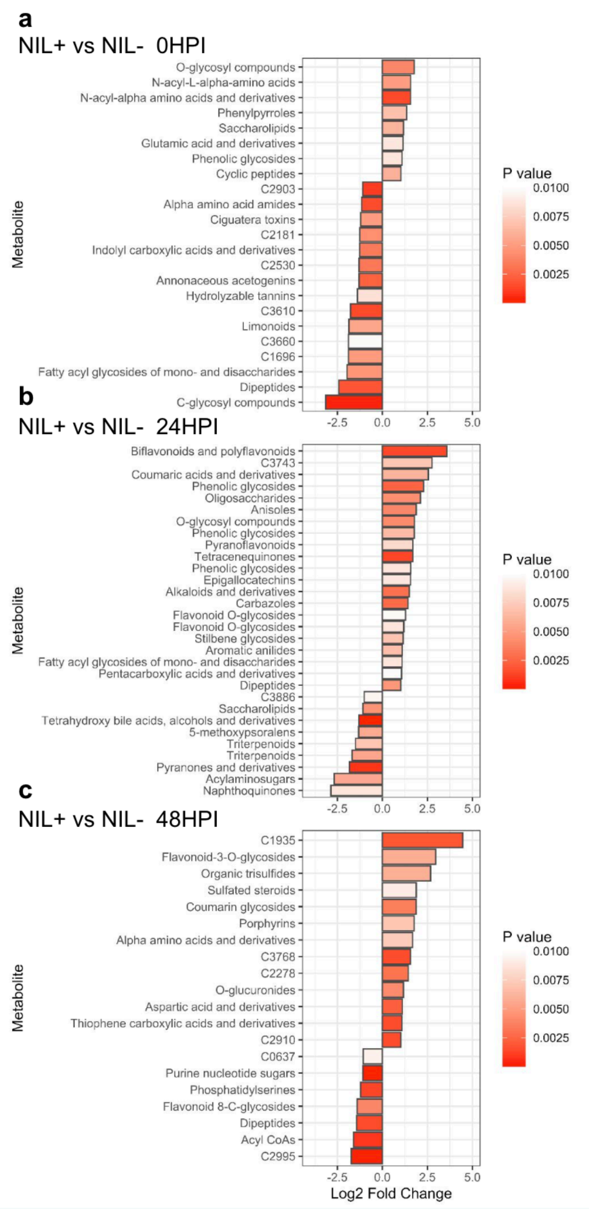


**Figure S2: Differential expression of sorghum metabolites by genotype for each level of treatment.** Log2 fold change is in relation to NIL+ at each timepoint. Analytes are labeled by their parent class, full metabolome annotations provided in Supplemental Data 2.

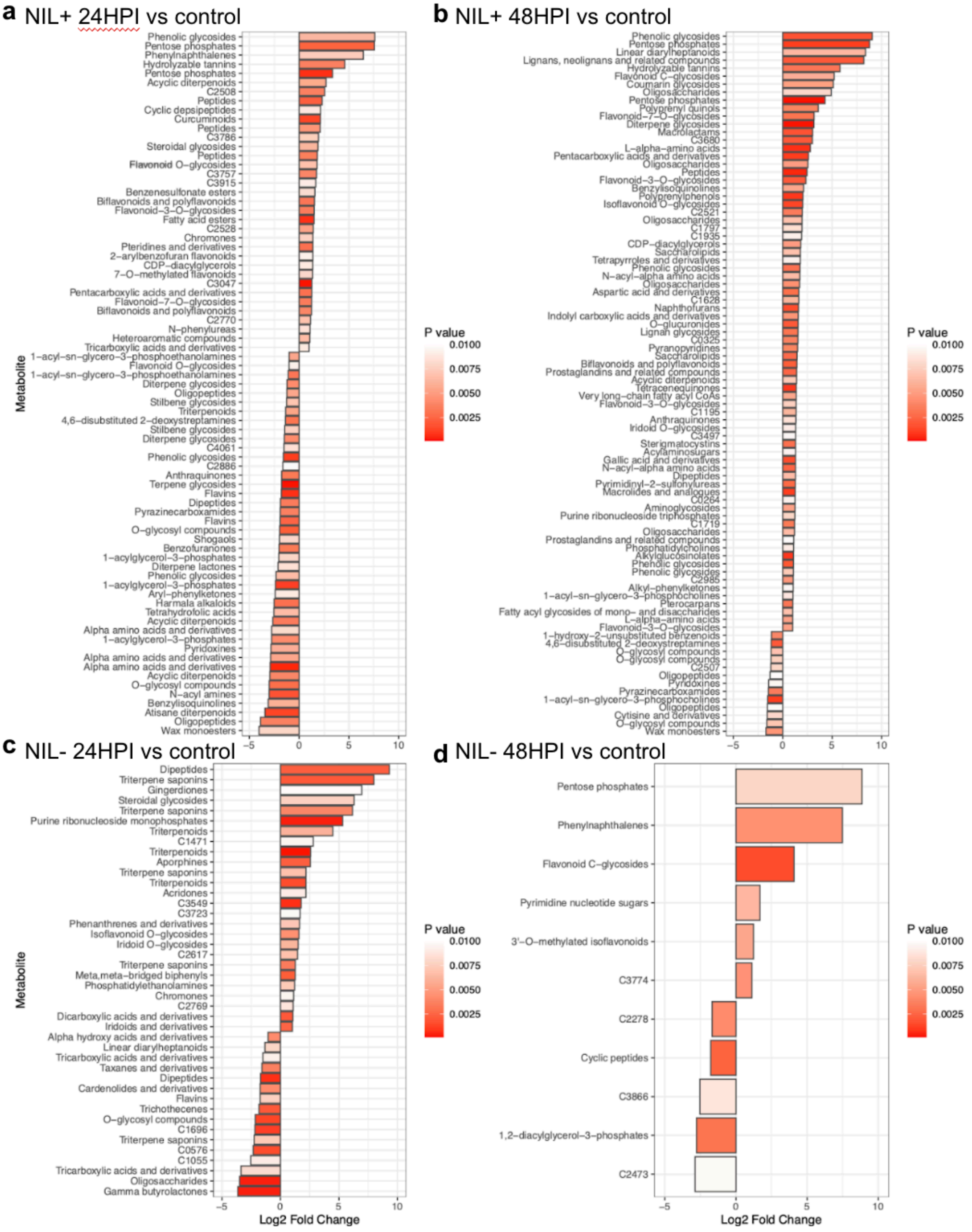


**Figure S3: Differential expression of sorghum metabolites in response to infestation.** Log2 fold change is in relation to NIL+ at each timepoint. Analytes are labeled by their parent class, full metabolome annotations provided in Supplemental Data 2.

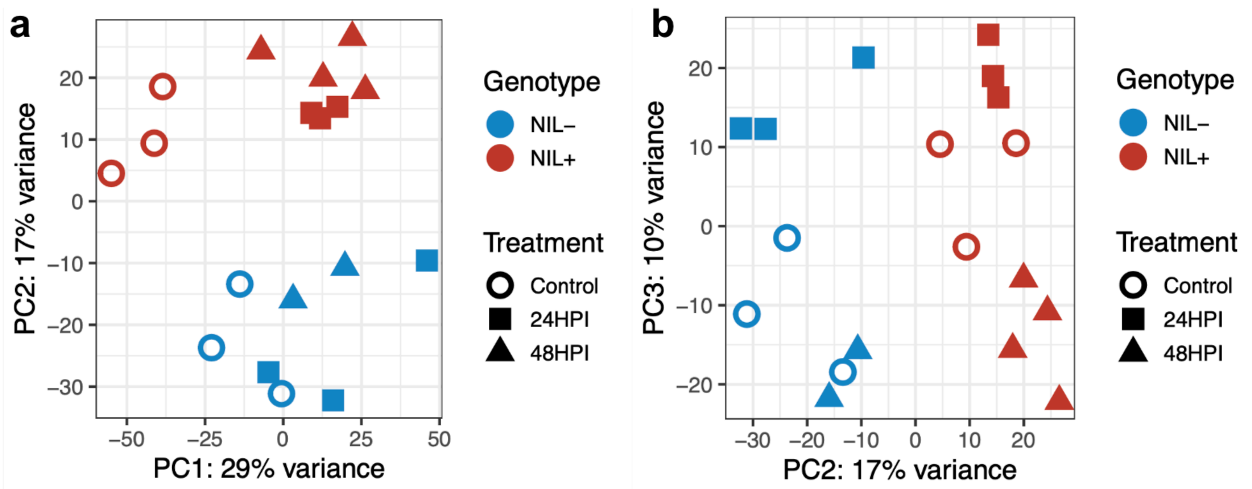


**Figure S4: Principal component analysis of sorghum transcriptomes (reference RTx430) using RMES1 near-isogenic lines.** a) PC1 vs PC2 as shown in main text. b) PC2 vs PC3. HPI, hours post infestation.

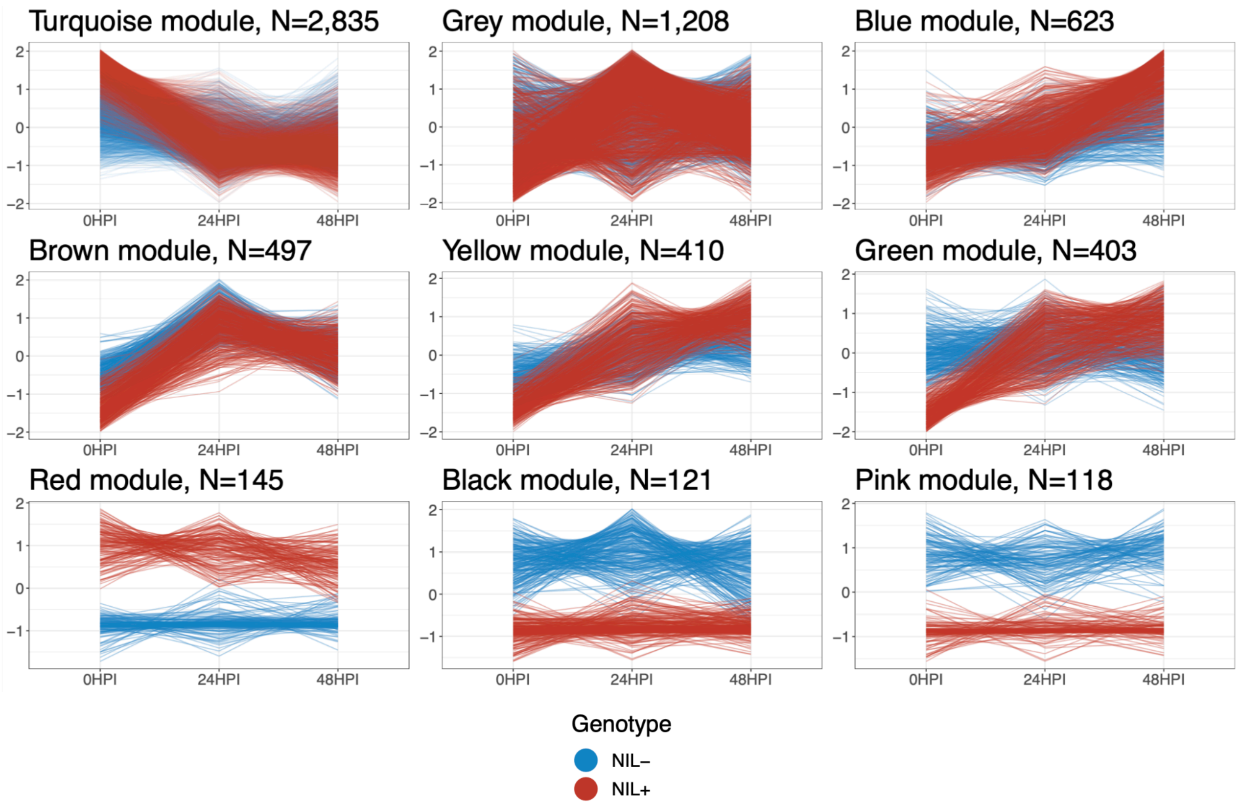


**Figure S5: Coexpression modules of sorghum genes (RTx430) in near-isogenic lines during aphid infestation.** Modules are labeled by their color i.e. MEturquoise is Turquoise module. Number of genes in each module is indicated above. HPI, hours post infestation.

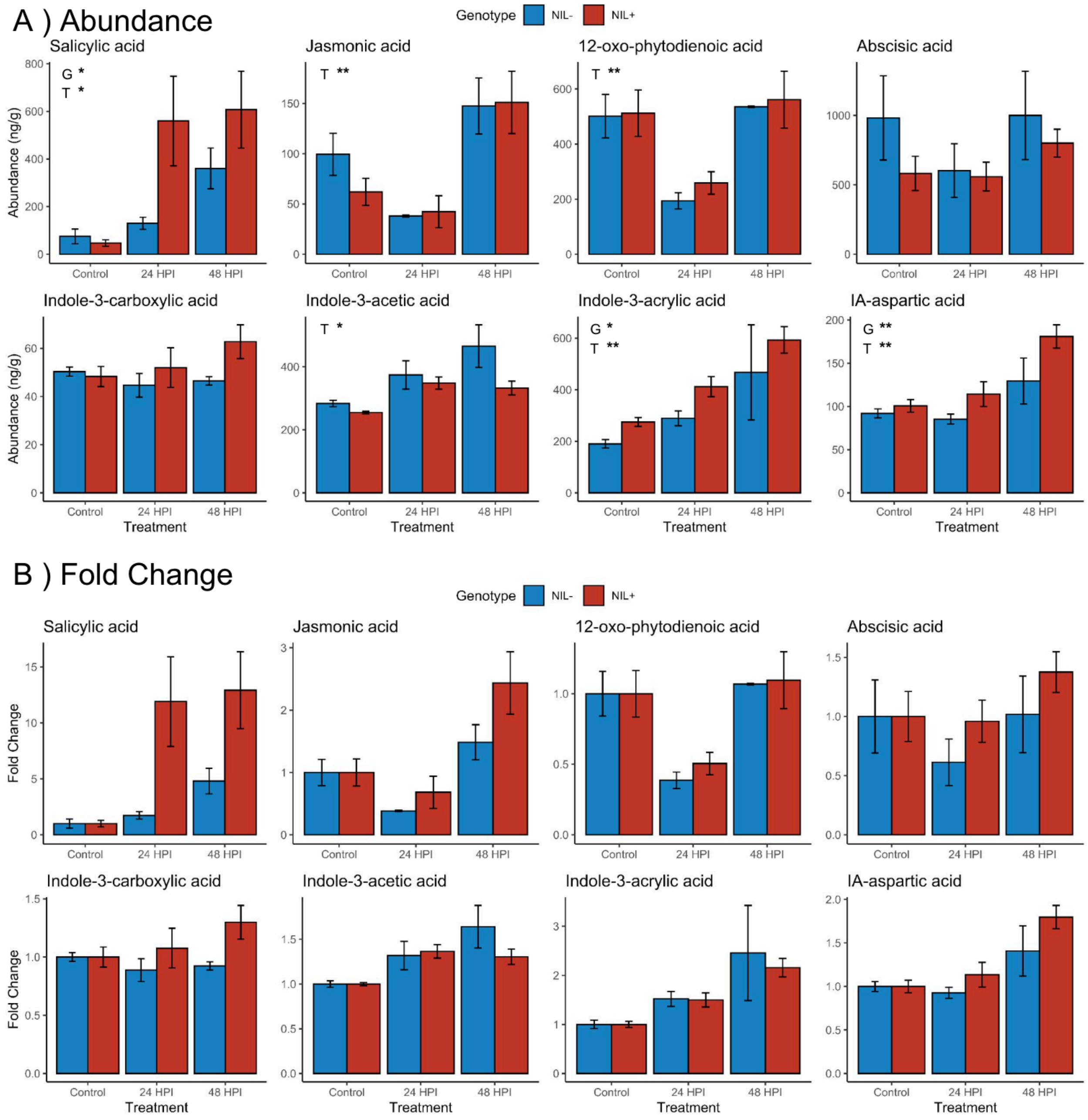


**Figure S6: Phytohormone abundance (a) and fold change (b) in sorghum near-isogenic lines during aphid infestation.** Genotype (G) and treatment (T) effect were determined using ANOVA, * = P-value < 0.05, ** = P-value < 0.01. HPI, hours post infestation.

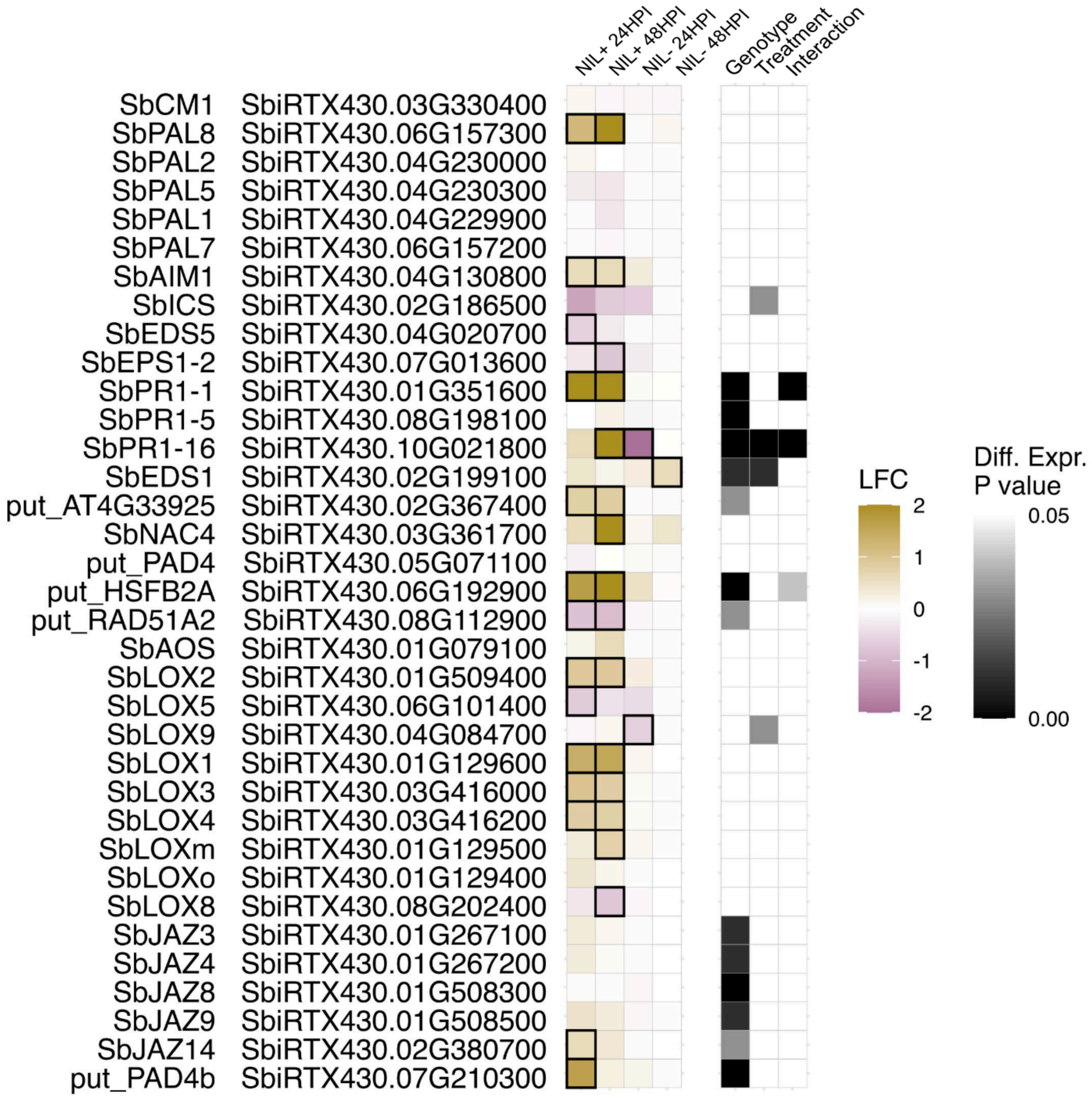


**Figure S7: Differential expression of sorghum genes (reference RTx430) in phytohormone and defense pathways in response to infestation.** Log fold change (LFC) is in relation to uninfested controls of the same genotype.

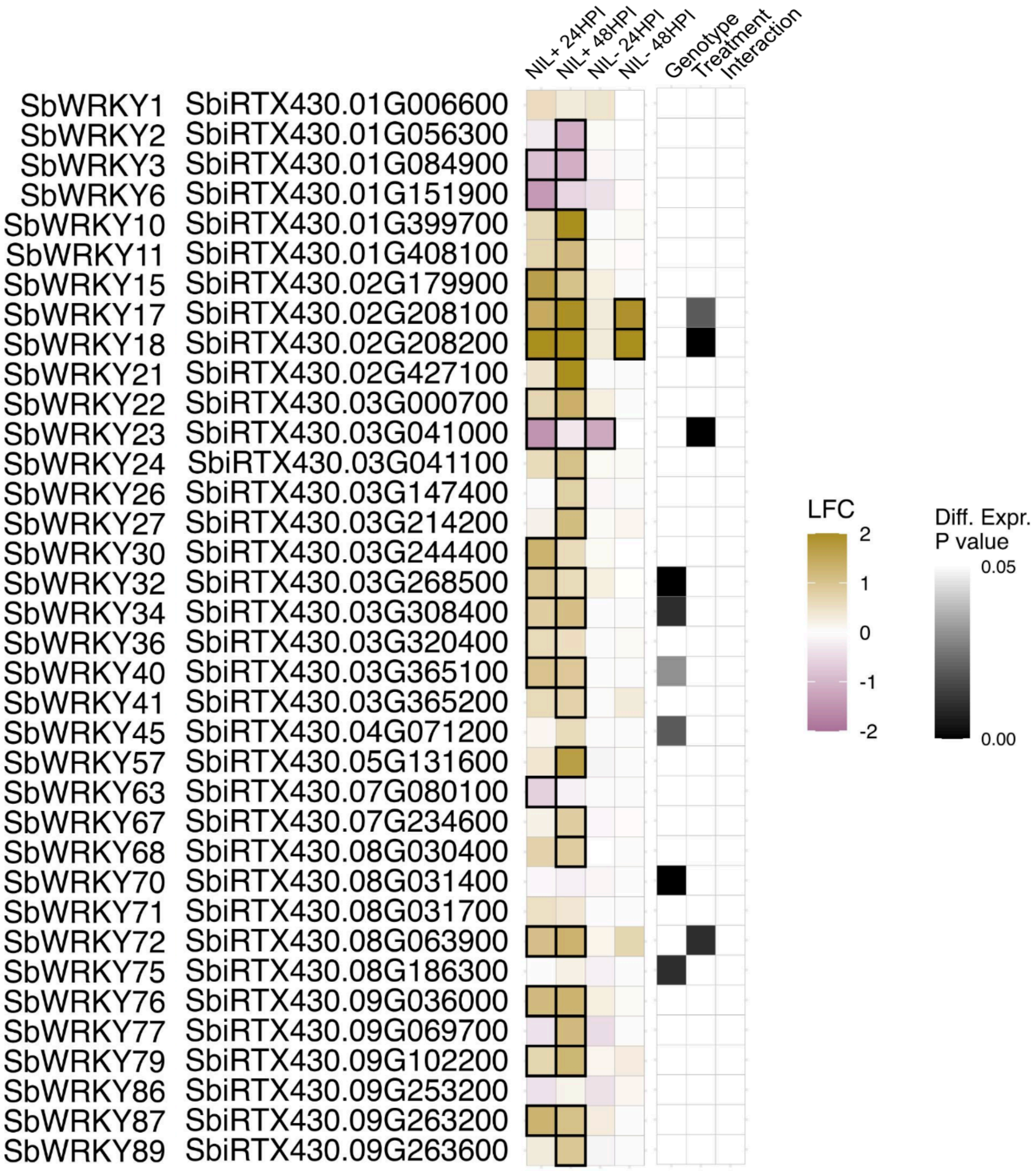


**Figure S8: Differential expression of WRKY transcription factor sorghum genes (reference RTx430) in response to infestation.** Log fold change (LFC) is in relation to uninfested controls of the same genotype.

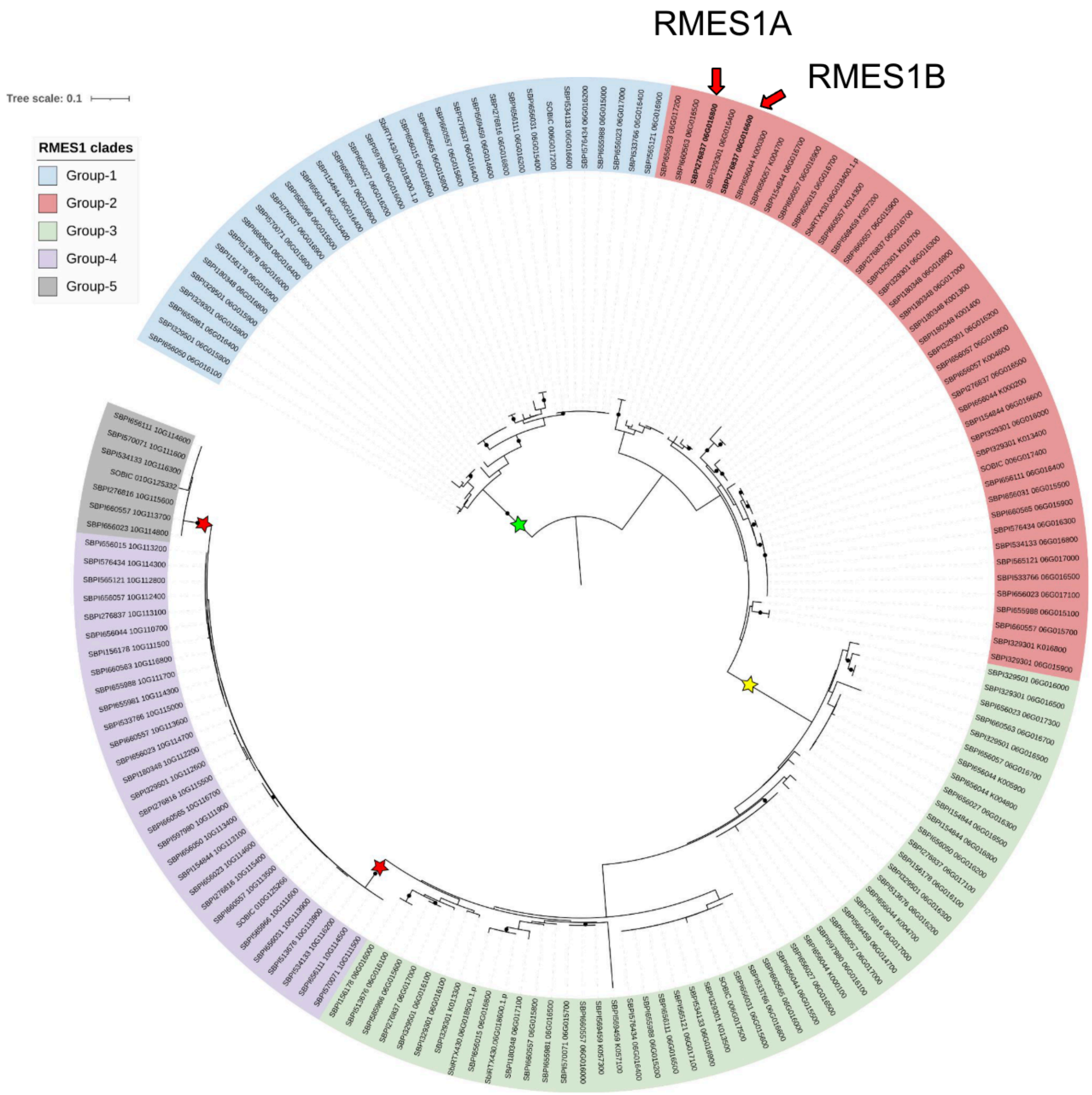


**Figure S9: Phylogenetic tree of global orthologs of RMES1 in sorghum pan-proteome.** Clades (group-1 to group-5) were determined by manual inspection of long, well-supported branches labeled by stars.

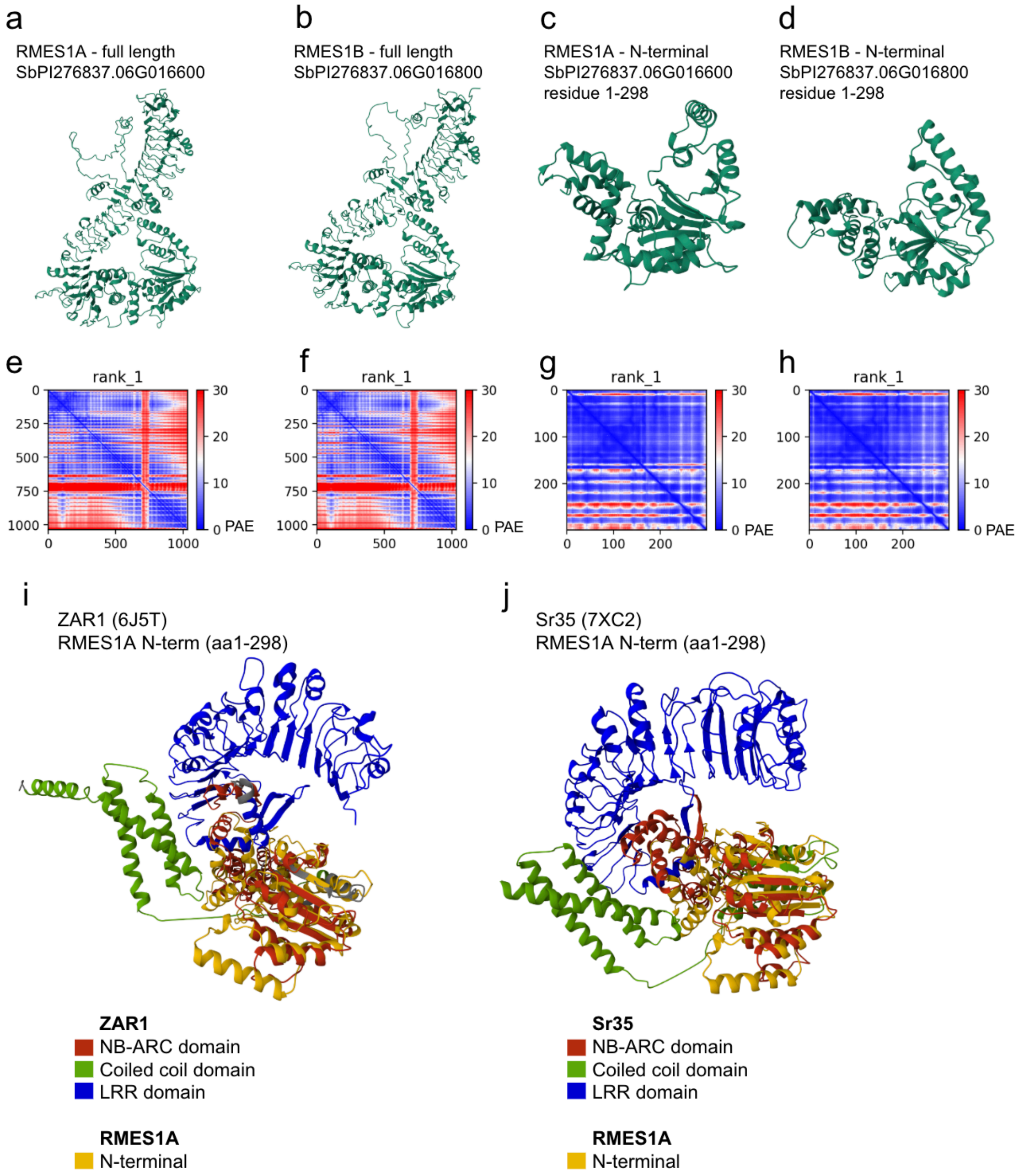


**Figure S10: AlphaFold structures of RMES1A, RMES1B, and alignment with ZAR1 and Sr35.** a-b) AlphaFold predicted structures of full-length and N-terminal sequence of RMES1A and RMES1B. e-f) Predicted aligned error (PAE) of AlphaFold hypotheses from panels a-d. i-j) FoldSeek alignment of RMES1A N-terminal (panel c) to ZAR1 and SR35 NBS domain.

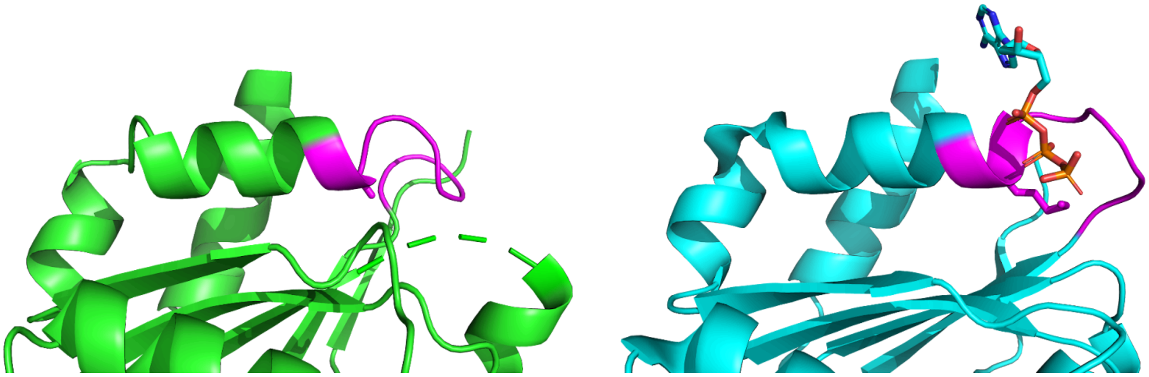

**Figure S11: AlphaFold predicted structure of RMES1A and ZAR1 P-loop sequence.** P-loop sequence (GxxxxGK[T/S], magenta) of the RMES1A NBS-like region (green) and ZAR1 NBS domain (blue). The sidechain of RMES1 A46 and ZAR1 K195 are shown. An ATP nucleotide shown in relation to the canonical ZAR1 P-loop.

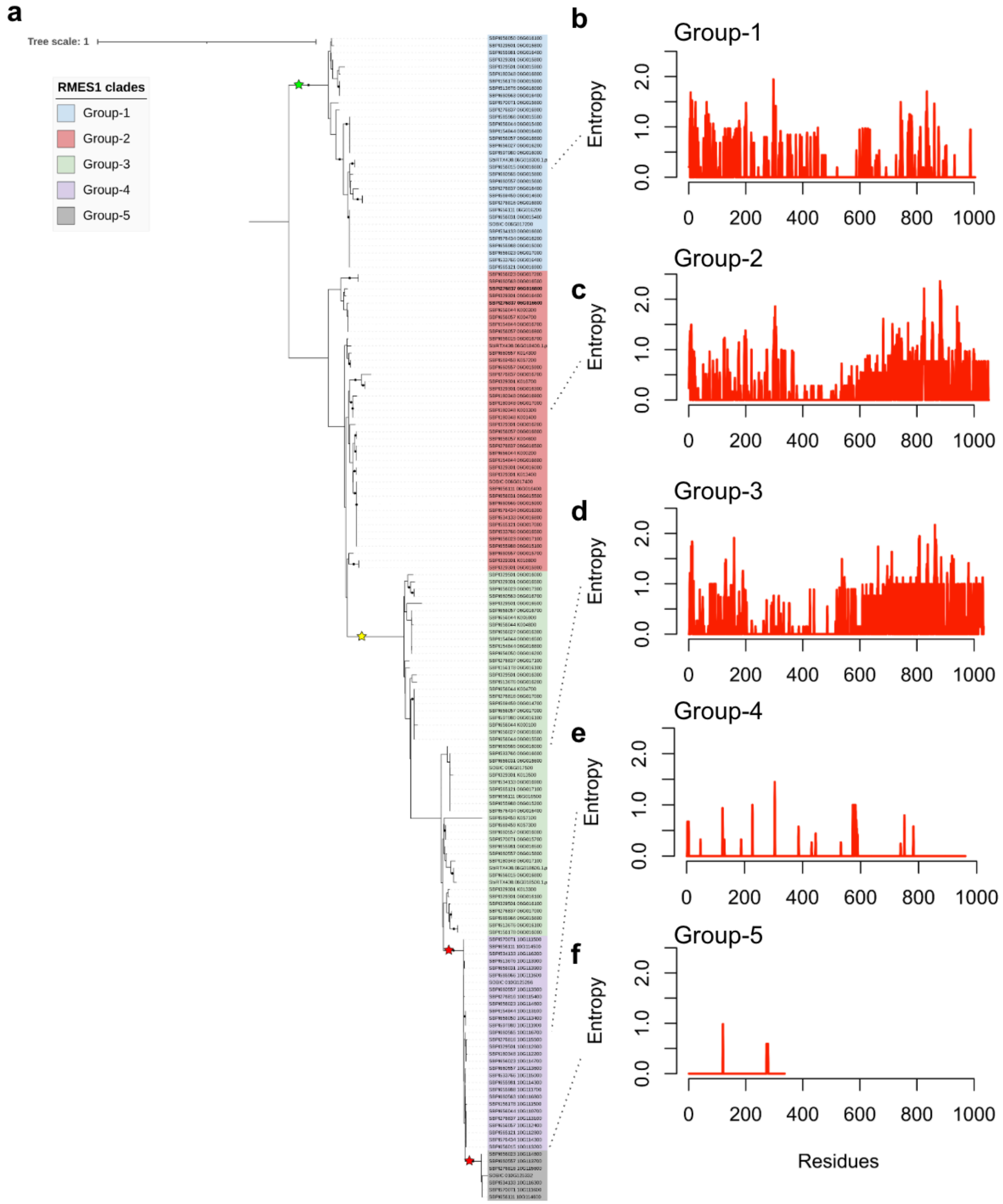


**Figure S12: Highly variable residues of RMES1 sorghum orthologs.** a) A linear view of the phylogenetic tree from Fig. S9 used to determine allelic series by clade. b-f) Shannon's entropy of residue positions. An entropy score of 0 indicates invariable positions, increasing scores indicate higher variability in amino acid identity.

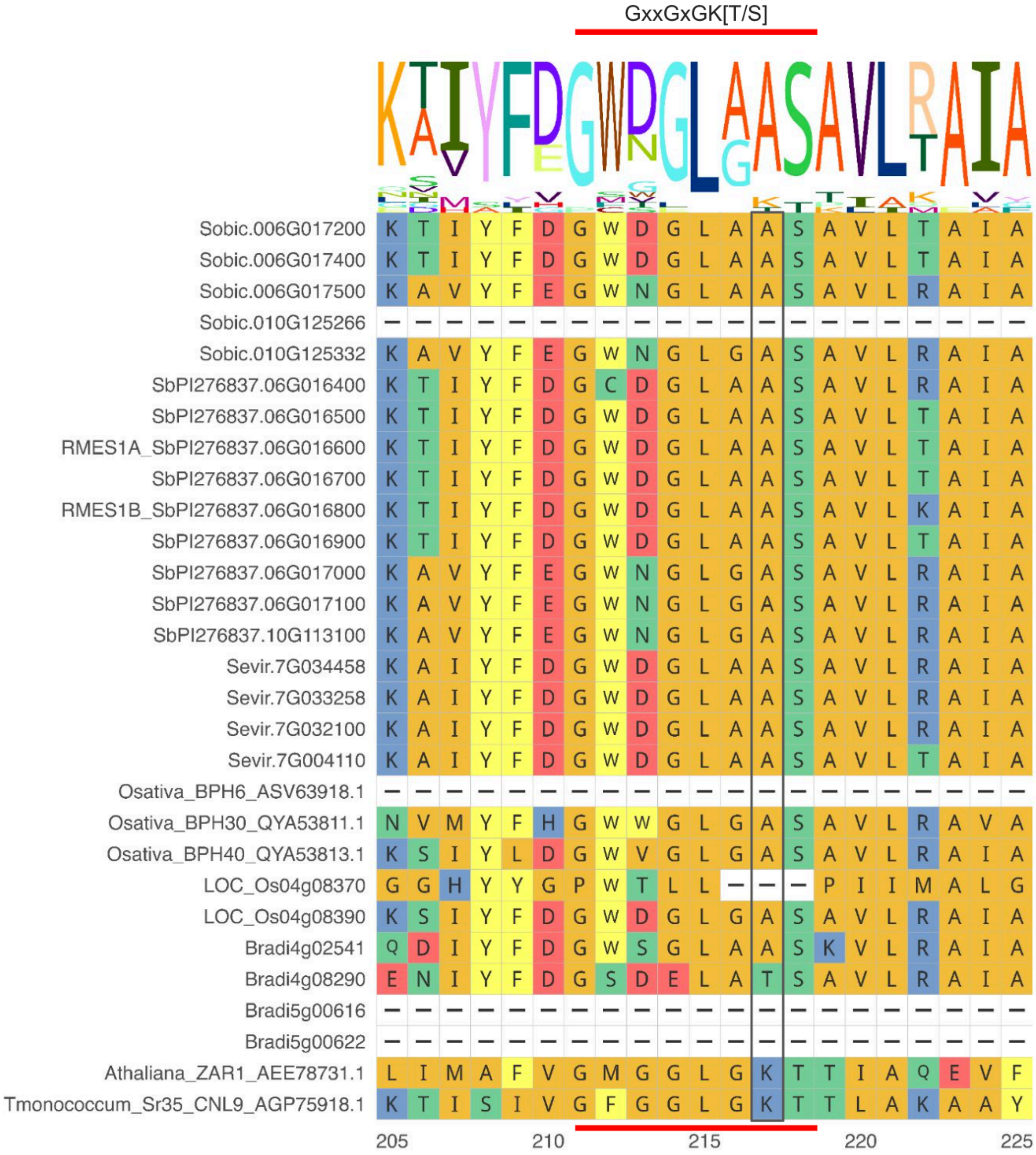


**Figure S13: Multiple sequence alignment of P-loop sequence for RMES1 orthologs across the super-pangenome of Poaceae.** The canonical P-loop motif is indicated above the alignment and the K/A variant expected to impair function is outlined.

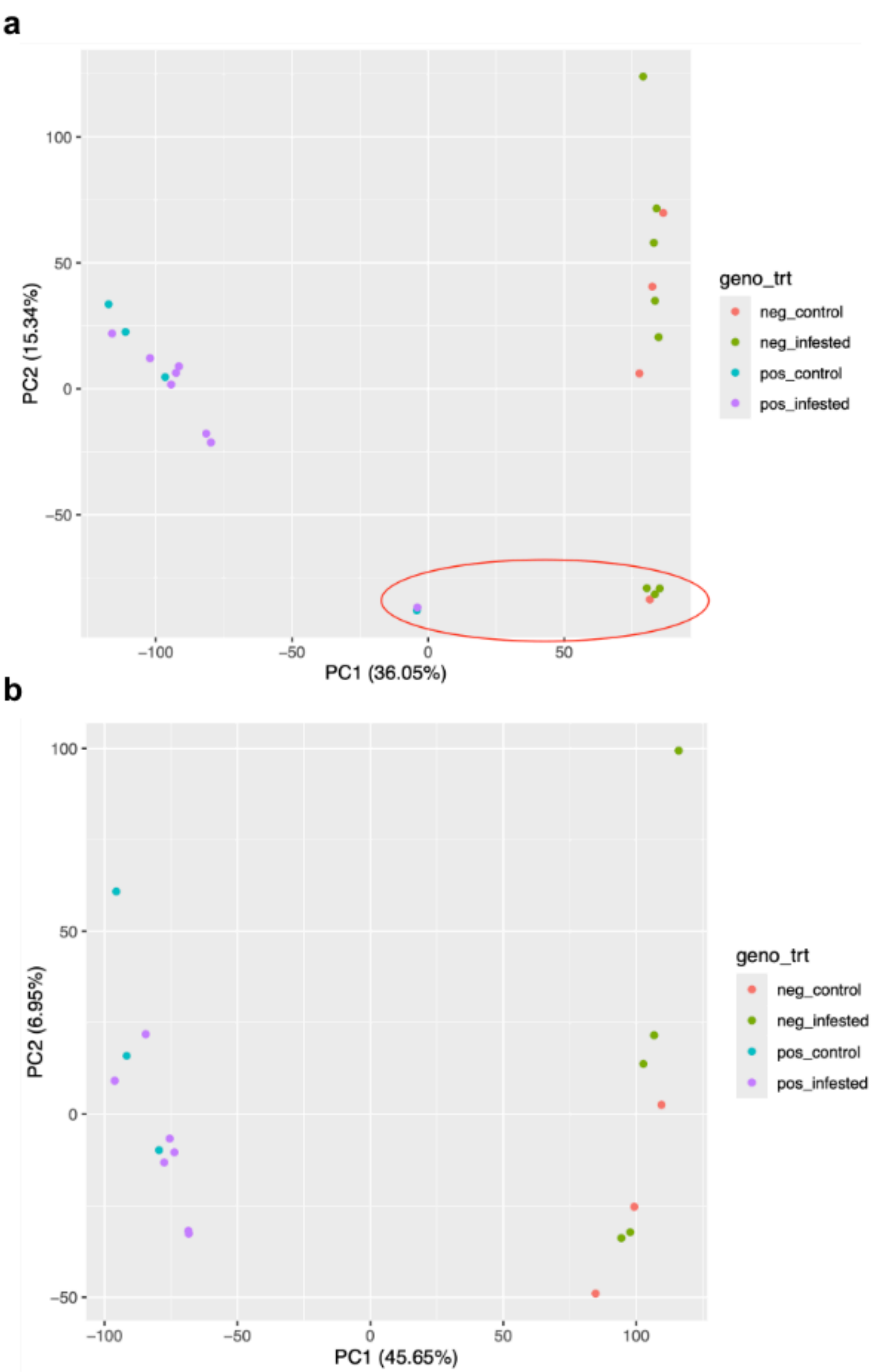


**Figure S14: Principal component analysis (PCA) of SNPs in sorghum near-isogenic lines.** a) SNPs of all 24 samples. Six samples with unexpected genomic makeup were identified and removed. b) PCA of remaining 18 samples which were used for transcriptome and metabolome analyses.

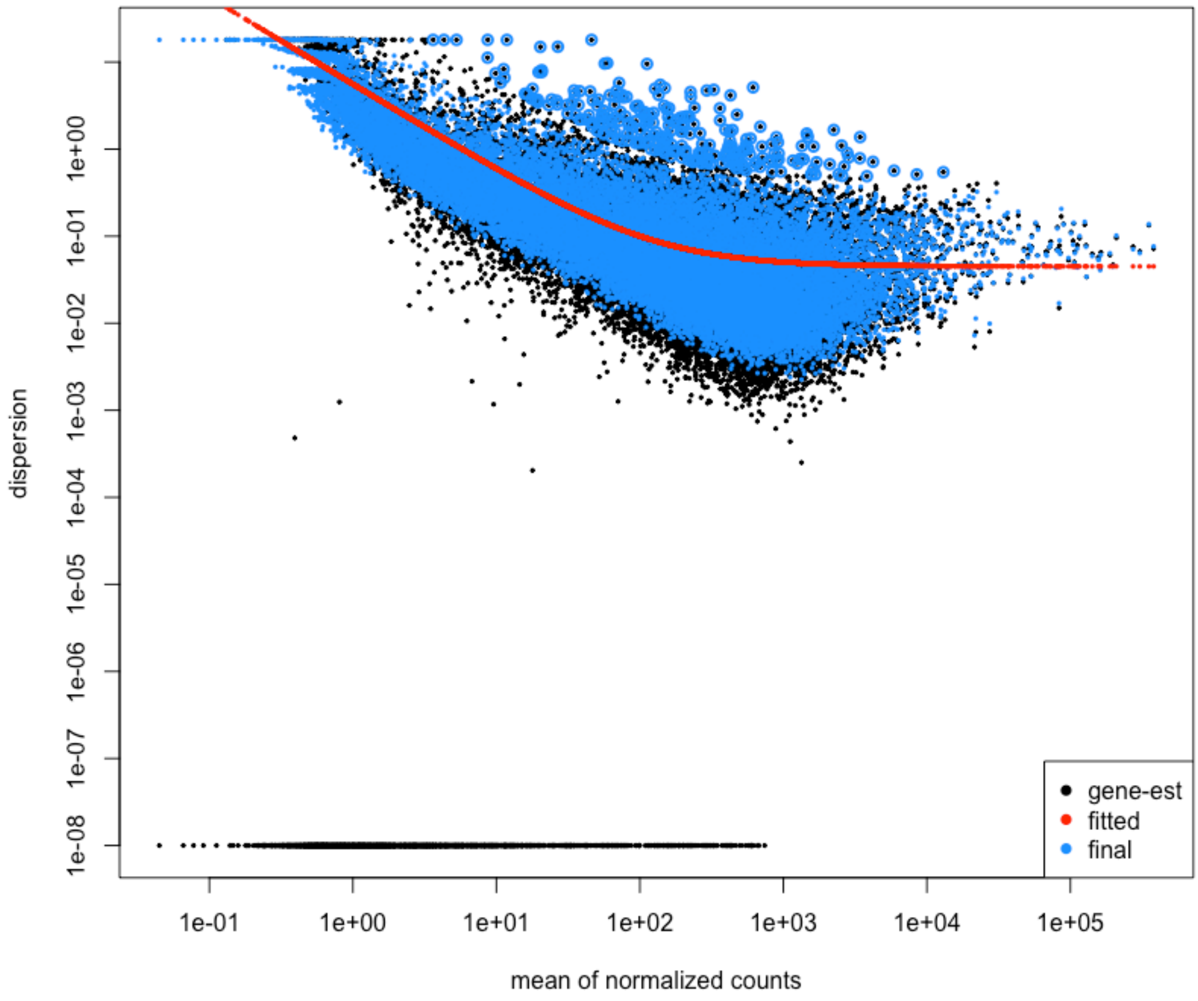


**Figure S15: Dispersion estimate plot of sorghum differentially expressed genes (reference RTx430) determined using DESeq2.** Red fitted line indicates dispersion across genes, black dots are raw dispersion estimates per gene, and blue circles denote Bayesian shrunken dispersions. A decreasing fitted line that plateaus at higher expression and shrunken values following the fitted line indicates the DESeq2 model is appropriate for identifying differentially expressed genes.
